# PathMHC: a workflow to selectively target pathogen-derived MHC peptides in discovery immunopeptidomics experiments for vaccine target identification

**DOI:** 10.1101/2024.09.11.612454

**Authors:** Owen Leddy, Yuko Yuki, Mary Carrington, Bryan D. Bryson, Forest M. White

## Abstract

Vaccine-elicited T cell responses can contribute to immune protection against emerging infectious disease risks such as antimicrobials-resistant (AMR) microbial pathogens and viruses with pandemic potential, but rapidly identifying appropriate targets for T cell priming vaccines remains challenging. Mass spectrometry (MS) analysis of peptides presented on major histocompatibility complexes (MHCs) can identify potential targets for protective T cell responses in a proteome-wide manner. However, pathogen-derived peptides are outnumbered by self peptides in the MHC repertoire and may be missed in untargeted MS analyses. Here we present a novel approach, termed PathMHC, that uses computational analysis of untargeted MS data followed by targeted MS to discover novel pathogen-derived MHC peptides more efficiently than untargeted methods alone. We applied this workflow to identify MHC peptides derived from multiple microbes, including potential vaccine targets presented on MHC-I by human dendritic cells infected with *Mycobacterium tuberculosis*. PathMHC will facilitate antigen discovery campaigns for vaccine development.

## Introduction

Antimicrobial resistant (AMR) bacterial, fungal, and parasitic infections pose a substantial threat to global public health, causing millions of deaths per year.^1^ The impact of AMR infections is expected to further increase by 2050, reaching USD 0.33-1.2 trillion per year in healthcare expenditures.^2^ Effective vaccines against AMR pathogens could mitigate these impacts and limit further opportunities for resistance to develop by preventing infections that would otherwise need to be treated with antimicrobials. T cell responses are essential and in some cases sufficient for immune protection against many common AMR pathogens and other globally significant microbial infections, including *Mycobacterium tuberculosis*^3^, uropathogenic *E. coli*^4^, *Salmonella* sp.^5^, *Staphylococcus aureus*^6^, *Streptococcus pneumoniae*^7^, *Klebsiella pneumoniae*^8,9^, *Trypanosoma cruzi*^10^, and others. In pathogens with multiple serotypes, T cell responses can confer broad-spectrum protection against distinct strains that lack antibody cross-reactivity.^7,8^ However, it is often not known which proteins are optimal targets for T cell priming vaccines. Methods to identify appropriate targets can therefore contribute to accelerating vaccine development against microbial pathogens.

T cell responses also contribute to immune protection against classes of viruses with pandemic potential, such as coronaviruses^11–13^ and influenza viruses^14,15^, and often target epitopes conserved in variants that evade antibody neutralization.^11,12,14^ The U.S. Biomedical Advanced Research and Development Authority (BARDA) has made establishing the capability to develop and deploy a vaccine within 100 days of a novel infectious disease outbreak a key strategic goal.^16^ Methods that can rapidly identify potential targets of protective T cell responses could contribute to achieving this goal and enhancing future outbreak preparedness.

Mass spectrometry analysis of MHC-bound peptides (termed immunopeptidomics) can directly identify which pathogen-derived antigens are presented on MHCs in infected cells and/or professional antigen-presenting cells and could serve as targets for protective T cell responses.^17,18^ Immunopeptidomics can have particular advantages as a vaccine target discovery approach in the context of microbial infections, since microbial pathogens may have thousands of protein-coding genes that can be challenging or costly to comprehensively screen for T cell reactivity. By identifying MHC peptides in an untargeted manner, immunopeptidomics allows proteome-wide discovery of potential antigenic targets, without an *a priori* bias toward any particular subset of the pathogen proteome. Immunopeptidomics can serve as a faster, less costly, and higher-throughput way to identify candidate human T cell antigens than approaches that rely on directly measuring T cell reactivity, which often require recruiting cohorts of human subjects with prior exposure to the pathogen of interest and may involve synthesizing large libraries of peptides.^19^

Pathogen-derived peptides can be outnumbered by self peptides in the MHC repertoire of infected cells by as much as 1000 to 1 and may be missed in untargeted immunopeptidome analyses performed using data-dependent analysis DDA.^20–22^ In DDA analyses, instrument control software selects precursor ions for fragmentation and MS/MS spectrum acquisition in real time, generally prioritizing more abundant precursor ions first with no prior knowledge of which peptides are pathogen-derived. Most instrument time during an analysis may be dedicated to MS/MS acquisition of self peptides, which are not candidate vaccine targets. In principle, targeted MS methods could selectively and sensitively identify pathogen-derived peptides while ignoring self peptides, but these methods require prior knowledge of peptides of interest and therefore cannot be used for novel target discovery.

Here, we describe a novel immunopeptidomics approach (termed “PathMHC”) that facilitates discovery of pathogen-derived MHC peptides by combining computational analysis of untargeted MS data with follow-up targeted MS analyses to identify previously missed pathogen-derived peptides. We first benchmark PathMHC against a comparable approach based on untargeted MS alone, using the non-pathogenic mycobacterium *M. smegmatis* as a model system. We next demonstrate that the PathMHC computational pipeline generalizes across infectious disease areas by applying it to publicly available immunopeptidomics datasets. Finally, we apply PathMHC to analyze the MHC class I (MHC-I) repertoire of human dendritic cells infected with *Mycobacterium tuberculosis* (*Mtb*), a globally important pathogen against which T cell-mediated immunity is an essential component of protective immune responses.^3^

## Results

### Combining computational enrichment with targeted MS facilitates identification of microbe-derived MHC peptides

The PathMHC workflow (Figure 1 a) requires MHC peptide samples isolated from cells infected with a pathogen of interest and from mock-infected control cells. First, each sample is divided in two equal parts, and one half of each sample (infected and mock-infected) is analyzed by untargeted MS (DDA). Second, the untargeted MS data are analyzed using a computational pipeline that finds precursor ions that are specific to infected cells and absent in the mock-infected control, excluding those already identified in the untargeted analysis (see Methods). Finally, this set of putatively infection-specific precursor ions is used as an inclusion list for a targeted MS analysis of the remaining half of each sample using parallel reaction monitoring (PRM) to identify pathogen-derived peptides previously missed in the untargeted analysis.

**Figure 1.**
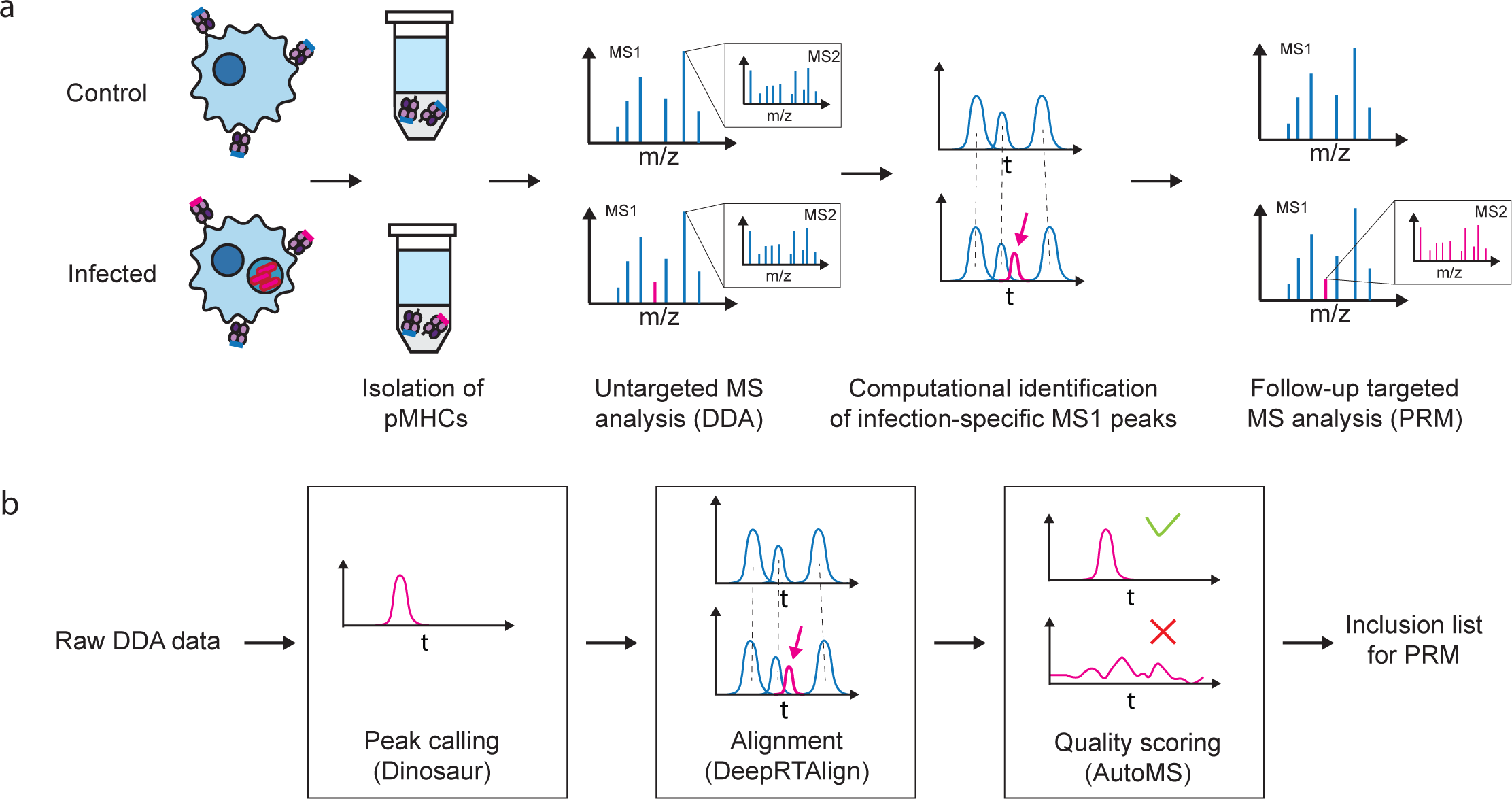
Overview of the PathMHC workflow. a) A schematic representation of the PathMHC workflow, including MHC peptide isolation from infected cells and mock-infected control cells, untargeted DDA analysis, computational identification of infection-specific precursor ions, and targeted follow-up PRM analyses. b) A schematic representation of the steps of the PathMHC computational pipeline, including peak calling using Dinosaur,^23^ chromatographic alignment using DeepRTalign,^24^ and peak quality filtering using AutoMS.^25^

The PathMHC computational pipeline comprises three steps (Figure 1 b): First, LC-MS features (i.e., precursor ions) are annotated using the feature detection algorithm Dinosaur.^23^ Second, the LC-MS features are aligned across the two samples using DeepRTAlign,^24^ and features in the infected sample that are not paired with any feature in the mock-infected control are selected for further analysis. Finally, LC-MS features that have a charge state and retention time consistent with MHC peptides are scored for quality (including signal-to-noise ratio) using the autoencoder-based scoring algorithm AutoMS.^25^ Peaks that meet scoring thresholds are added to the inclusion list for the follow-up targeted analysis with PRM.

To benchmark the PathMHC approach, we used *Mycobacterium smegmatis* (*Msmeg*), a non-pathogenic mycobacterium. We incubated primary human monocyte-derived dendritic cells (hMDCs) with *Msmeg* at a multiplicity of infection (MOI) of 10 for 2 hours to allow phagocytosis of the bacteria, harvested and lysed the cells 24 hours after phagocytosis, and isolated MHC-II-associated peptides (Figure 2 a).

**Figure 2.**
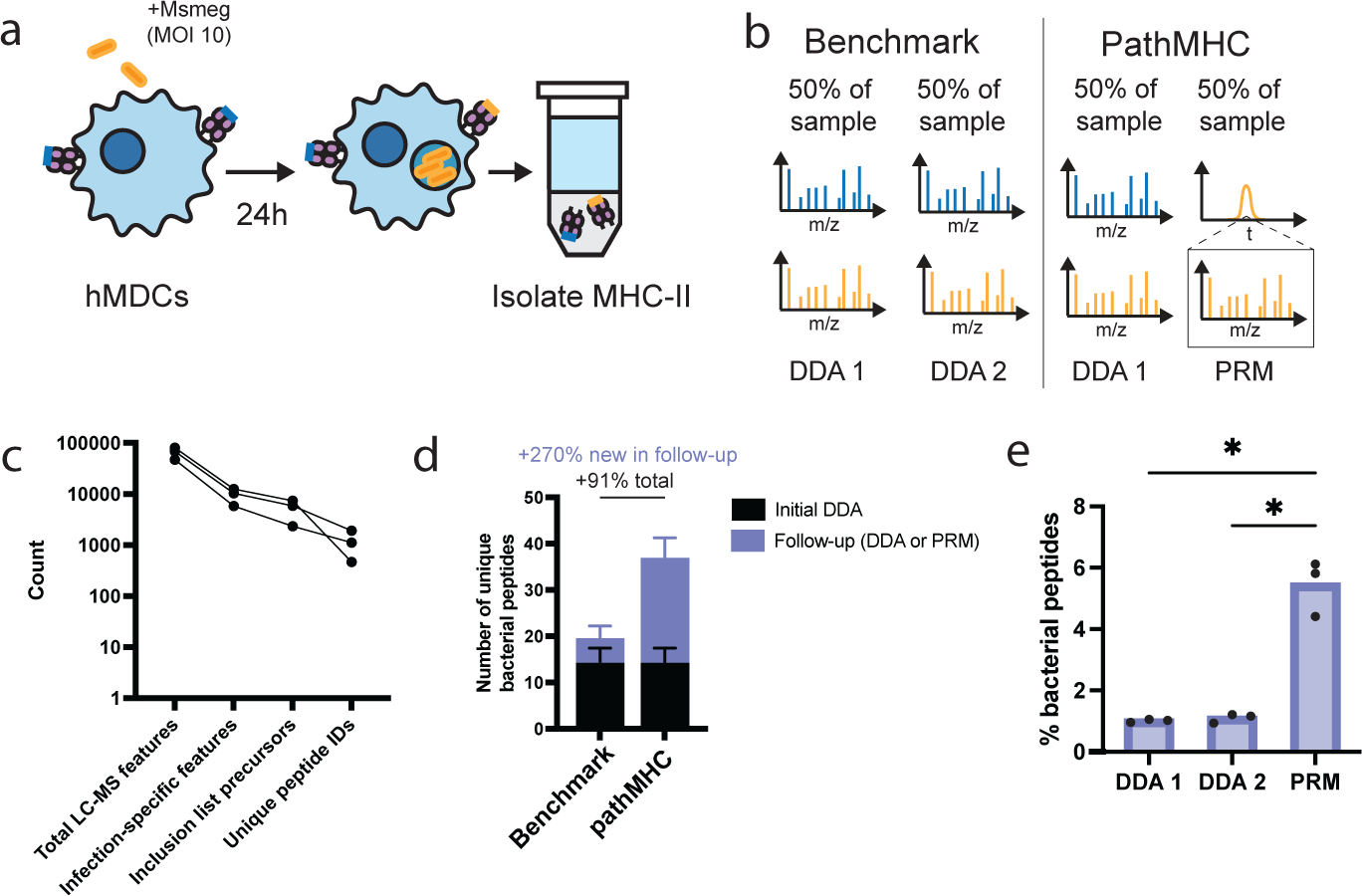
PathMHC identifies more bacterial peptides presented by professional antigen presenting cells than does DDA alone in a model system. a) Schematic representation of the model system used to benchmark PathMHC. Human monocyte-derived dendritic cells (hMDCs) were allowed to phagocytose *Msmeg* at an MOI of 10 for 2 hours, and 24 hours later MHC-II peptides were isolated for MS analysis. b) Outline of the benchmark DDA-only workflow and the corresponding PathMHC workflow. c) Number of LC-MS (precursor ion) features identified by Dinosaur, precursors with charge state +2 or +3 identified as specific to infected cells by DeepRTalign (“infection-specific features”), precursors selected for inclusion list, and final number of peptide IDs obtained by PRM, for n = 3 PathMHC analyses of hMDCs that phagocytosed *Msmeg*. d) Number of unique bacterial peptides identified in the initial DDA analysis (black bar) and the number of new unique bacterial peptides identified in the follow-up DDA or PRM analysis (lavender bar) for the benchmark or PathMHC workflow respectively. Bars represent the mean ± standard deviation of n = 3 replicates using cells from different healthy donors. e) The percentage of peptides derived from *Msmeg* in each DDA or PRM analysis, showing enrichment of bacterial peptides in PathMHC PRM follow-up analyses for n = 3 replicates using cells from different healthy donors (* p < 0.05, one-way paired ANOVA).

In early tests of prototype PathMHC workflows, injecting 100% of an MHC-II peptide sample in a single DDA MS analysis identified fewer bacterial peptides than did two consecutive DDA analyses, each using 50% of the sample (Supplementary figure 1). The inherent variability in selection of peptides for MS/MS spectrum acquisition during DDA analyses means that the second analysis will capture some bacterial peptides not identified in the first, resulting in more IDs as long as the bacterial peptides are still above the limit of detection when using 50% of the sample. We therefore adopted two consecutive DDA analyses as the baseline method against which we would benchmark PathMHC (Figure 2 b).

An average of 65,293 LC-MS features (precursor ion peaks) were identified in analyses of MHC-II peptides from *Msmeg*-infected hMDCs, 14.4% of which were classified as specific to infected cells and absent in the corresponding mock-infected control by DeepRTalign and had an appropriate charge state (+2 or +3). After discarding precursor ion peaks that were already identified in the initial untargeted MS analysis or failed to meet AutoMS quality scoring thresholds, an average of 7.57% of total precursor ions were ultimately added to the inclusion list for targeted analysis by PRM (Figure 2 c).

The PathMHC workflow (one untargeted DDA analysis and one targeted PRM analysis) on average identified approximately 91% more bacterial peptides than did two consecutive untargeted DDA analyses (Figure 2 d). PathMHC PRM analyses detected, on average, approximately 270% more new bacterial peptides (i.e., peptides that were not detected in the initial DDA analysis) than did a second DDA analysis of the same sample. On average, 1.01% of peptides identified in DDA analyses were *Msmeg*-derived, whereas 5.45% of peptides identified in PRM analyses using inclusion lists generated by the PathMHC computational pipeline were *Msmeg-*derived, showing that the PathMHC pipeline successfully enriched for bacterial peptides (p < 0.05; Figure 2 e). These results show that PathMHC can increase the number of bacterial MHC peptides identified from a given sample, compared to DDA alone. In addition to our pipeline based on open-source tools, an analogous computational pipeline implemented in a commercial proteomics analysis program can also enrich for bacterial MHC-II peptides, giving users of our method a choice of multiple possible software implementations (Supplementary note 1; Supplementary figure 2).

### The PathMHC computational pipeline generalizes across pathogens

We next wanted to test whether our PathMHC computational pipeline generalizes across different infectious disease areas and mass spectrometry instrumentation. We therefore used the PathMHC computational pipeline to identify infection-specific precursor ions in published untargeted MS analyses of the MHC-I repertoire of the A549 human epithelial cell line infected with SARS-CoV-2.^21^ In this case, we could not know what the identities of all the selected precursors would have turned out to be if they had been targeted in a follow-up PRM analysis, so we instead analyzed the subset of precursors selected by the PathMHC pipeline that had already been identified in the untargeted MS analysis. SARS-CoV-2-derived peptides were enriched among the precursors selected by the PathMHC computational pipeline that had known identities, relative to the overall set of peptides identified (Figure 3 a). Self peptides selected by the PathMHC pipeline were enriched for proteins associated with interferon responses, which are known to be induced by SARS-CoV-2 infection^26^ (Figure 3 b). These results indicate that self peptides selected by the PathMHC computational pipeline are enriched for pathogen-derived peptides and peptides derived from proteins associated with the host response to infection.

**Figure 3.**
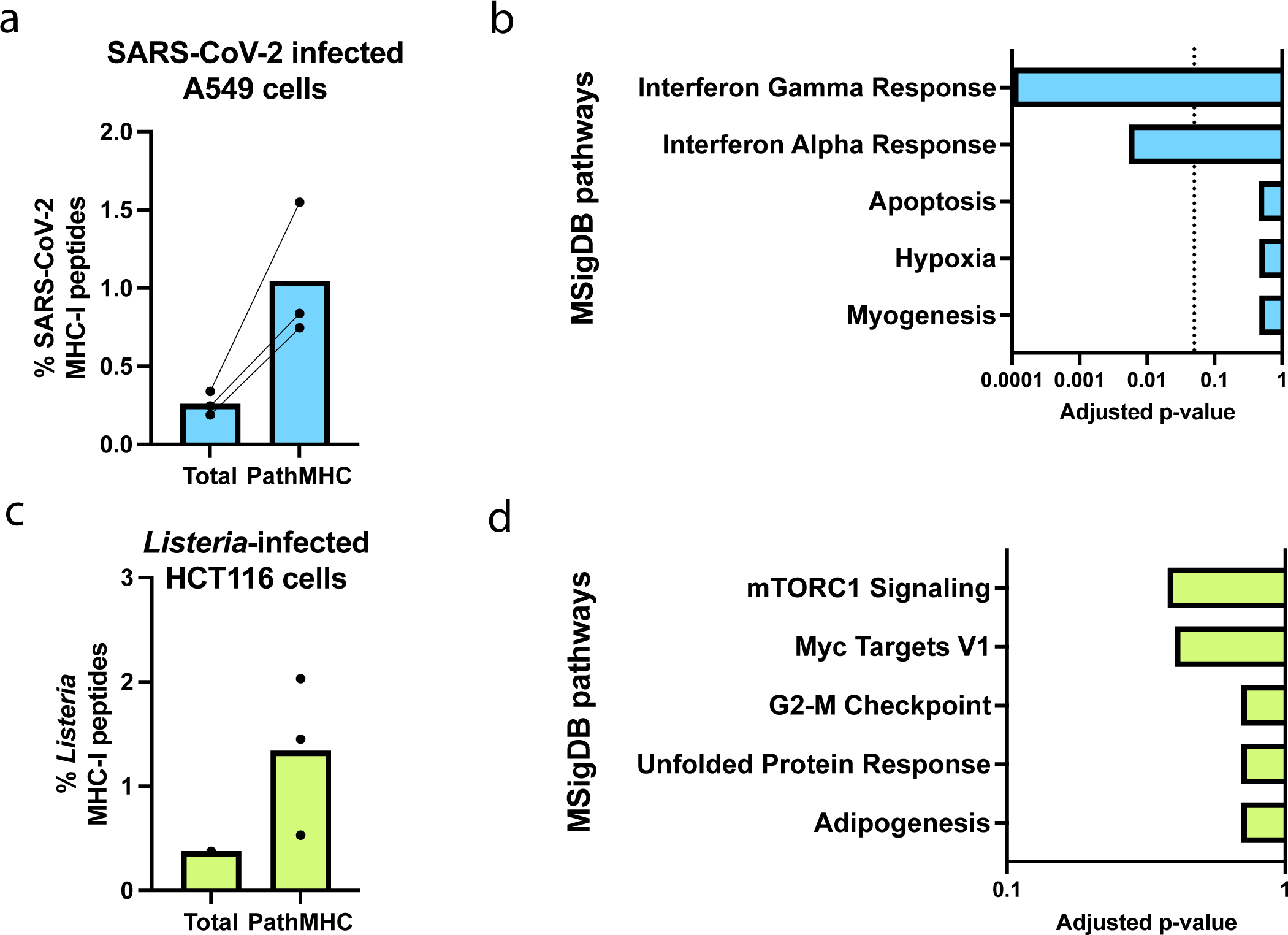
PathMHC enriches for pathogen-derived and infection-associated peptides in publicly available immunopeptidomics datasets. Percentage of pathogen-derived peptides in published DDA analyses of MHC-I peptides isolated from a) SARS-CoV-2-infected A549 cells (n = 3 fractions)^21^ and c) *Listeria monocytogenes*-infected HCT116 cells (n = 3 replicates),^20^ compared to the percentage of pathogen-derived peptides among identified precursors selected by the PathMHC computational pipeline (not excluding peptides identified by DDA; see Methods). Pathway enrichment analysis of precursors selected by the PathMHC computational pipeline that were identified as self peptides in b) SARS-CoV-2-infected A549 cells and d) *Listeria monocytogenes*-infected HCT116 cells. Enrichment analysis was performed using Enrichr^50^ against the MSigDB 2020 database,^52^ relative to the overall set of source proteins of peptides identified in the study as a background. The dashed line indicates the threshold for statistical significance after multiple hypothesis testing correction (Benjamini-Hochberg).

An analogous computational analysis of an MS dataset of MHC-I peptides isolated from the HCT116 human colon epithelial cell line infected with *Listeria monocytogenes* also enriched for *Listeria*-derived peptides relative to the overall proportion of *Listeria-*derived peptides identified in the study^20^ (Figure 3 c). (In this case, the study authors did not report IDs separately for each replicate.) Although not statistically significant after multiple hypothesis test correction, the mTORC1 signaling pathway was the most enriched among source proteins of self peptides selected by the PathMHC computational pipeline (Figure 3 d). *Listeria monocytogenes* has previously been shown to induce mTORC1 signaling to facilitate entry into host cells.^27^ The fact that the PathMHC computational pipeline enriches for SARS-CoV-2- and *Listeria*-derived MHC peptides in publicly available datasets suggests that PathMHC can help identify pathogen-derived MHC peptides across a range of infectious disease areas.

### PathMHC identifies potential TB vaccine targets presented on MHC-I

We next applied PathMHC to analyze the MHC-I repertoire of hMDCs infected with *Mtb* to identify potential TB vaccine targets. hMDCs were infected with *Mtb* at an MOI of 2.5 for 72 hours prior to isolation of MHC-I peptides (Figure 4 a). *Mtb*-derived peptides comprised 0.116% of MHC-I peptide IDs in infected hMDCs in DDA analyses and 0.156% on average in PRM analyses (Figure 4 b), suggesting a modest enrichment despite the low overall prevalence of *Mtb* peptides in this system. We obtained a total of 19 *Mtb*-derived peptide IDs (16 unique epitopes), including 4 identified in PRM analyses that had not been previously identified by DDA (2 unique epitopes) (Figure 4 c). Peptides identified by PathMHC PRM analyses that were previously missed in DDA analyses derived from EspC and the EsxJ family of proteins – EsxJ, EsxK, EsxP, and EsxW (abbreviated here as EsxJKPW).

**Figure 4.**
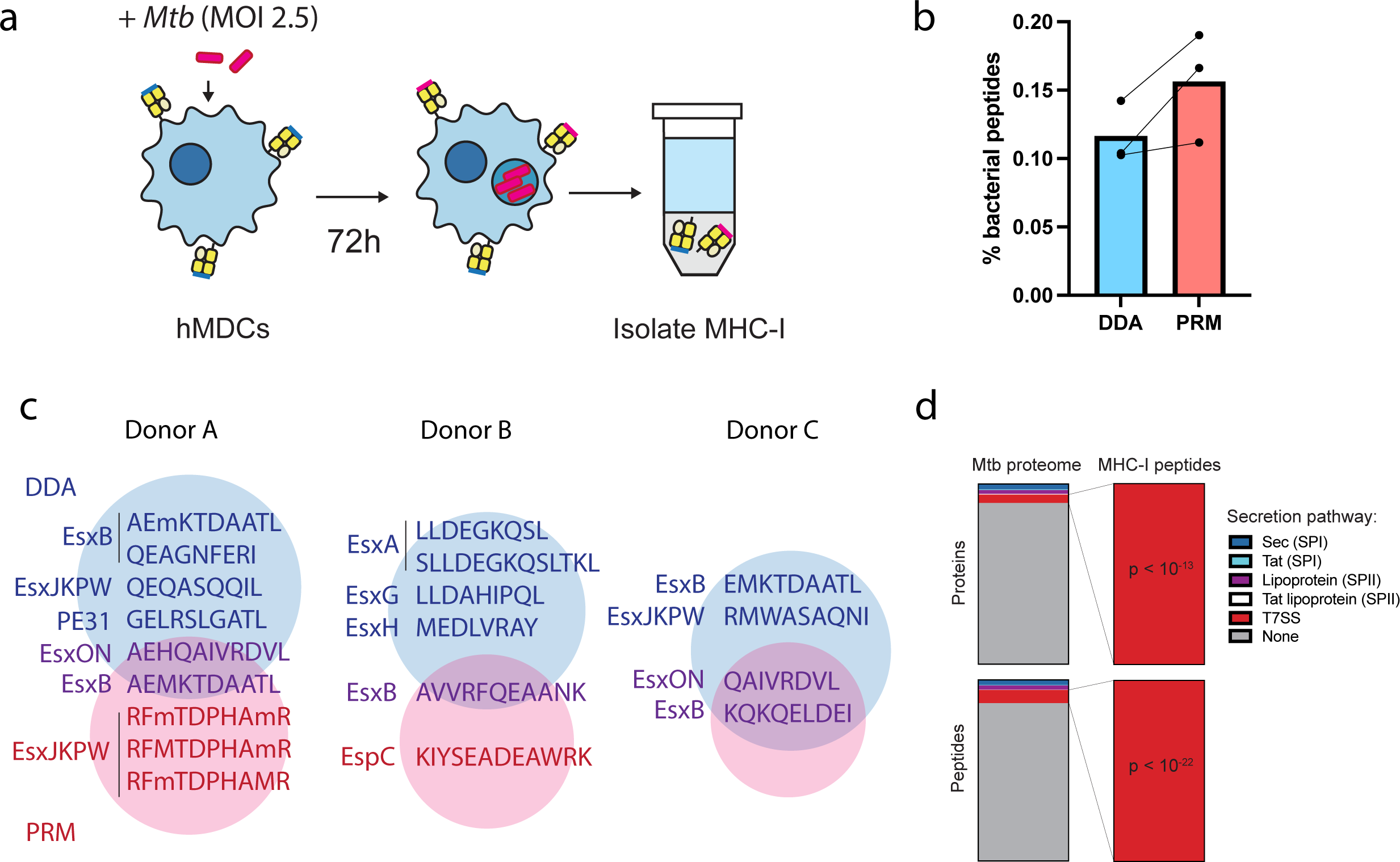
*Mtb*-infected hMDCs present peptides derived from T7SS substrates on MHC-I. a) Schematic representation of the *Mtb* dendritic cell infection model. hMDCs were allowed to phagocytose *Mtb* at an MOI of 2.5 for 4 hours, and 72 hours later MHC-I peptides were isolated for MS analysis. b) The proportion of *Mtb*-derived peptides in each initial DDA analysis and follow-up PRM analysis for each of n = 3 donors. c) *Mtb* peptide sequences detected in DDA analyses, PRM analyses, or both in the PathMHC workflow (right) and their source proteins (left) for each of three donors. Lowercase m = oxidized methionine. d) Enrichment analysis of *Mtb* protein secretion pathways among the detected MHC-I peptides relative to the encoded *Mtb* H37Rv proteome, by peptide (binomial test) and by source protein (hypergeometric test; see Methods).

We validated the *Mtb* MHC-I peptide IDs we obtained from PathMHC by SureQuant^22^ (Supplementary Figure 3) (a targeted MS method that compares biological MHC peptides with stable isotope labeled synthetic standards to demonstrate identical retention times and MS/MS spectra) along with 5 other epitopes that had been separately identified in DDA-only analysis of MHC-I peptides from hMDCs (Supplementary Figure 4). Unsupervised clustering of MHC-I peptides using Gibbs clustering^28,29^ resulted in clusters with sequence motifs that matched the known peptide binding preferences of HLA alleles expressed by each donor (Supplementary figure 5). Each *Mtb*-derived peptide was predicted to bind at least one class I human leukocyte antigen (HLA) allele expressed by the corresponding donor by the MHC peptide prediction algorithm NetMHCpan^30^ (Supplementary figure 6). These results increase our confidence that the peptides identified are authentic *Mtb* MHC-I epitopes.

The *Mtb-*derived MHC-I peptides presented by hMDCs exclusively derived from substrates of *Mtb*’s type VII secretion systems (T7SSs), a statistically significant enrichment relative to the encoded *Mtb* proteome (peptide enrichment: p < 10^-22^, binomial test with Bonferroni correction; source protein enrichment: p < 10^-13^, hypergeometric test with Bonferroni correction; Figure 4 d). T7SSs are specialized molecular machines that export proteins across the bacterial cell envelope and have roles in nutrient acquisition,^31^ outer membrane selective permeability,^32^ and virulence^33^. Four putative non-T7SS-derived *Mtb* peptides were selected for SureQuant validation but did not co-elute with the corresponding synthetic standards, indicating these IDs were incorrect (Supplementary table 2). We previously observed that *Mtb*-infected human monocyte-derived macrophages also predominantly present T7SS substrates on MHC-I.^22^ Our results here suggest that the determinants of *Mtb* MHC-I antigen processing and presentation may be similar among multiple classes of monocyte-derived phagocytes.^34,35^

PathMHC also targets self peptides that reflect the host cell response to *Mtb* infection. Self peptides uniquely detected by PathMHC PRM analyses of *Mtb*-infected hMDCs and not in DDA analyses were enriched for proteins associated with interferon responses, relative to self peptides identified by DDA (Figure 5 a). The type I interferon response is a dominant component of the myeloid cell response to *Mtb* infection.^36–38^ Many interferon-associated self peptides identified by PathMHC were only detectable in infected cells or highly upregulated during infection (Figure 5 b), whereas self MHC-I peptides identified in DDA analyses of mock-infected cells were variously upregulated, downregulated, or unaffected by *Mtb* infection (Supplementary Figure 7). These results show that PathMHC enriches for MHC-I peptides associated with *Mtb* infection.

**Figure 5.**
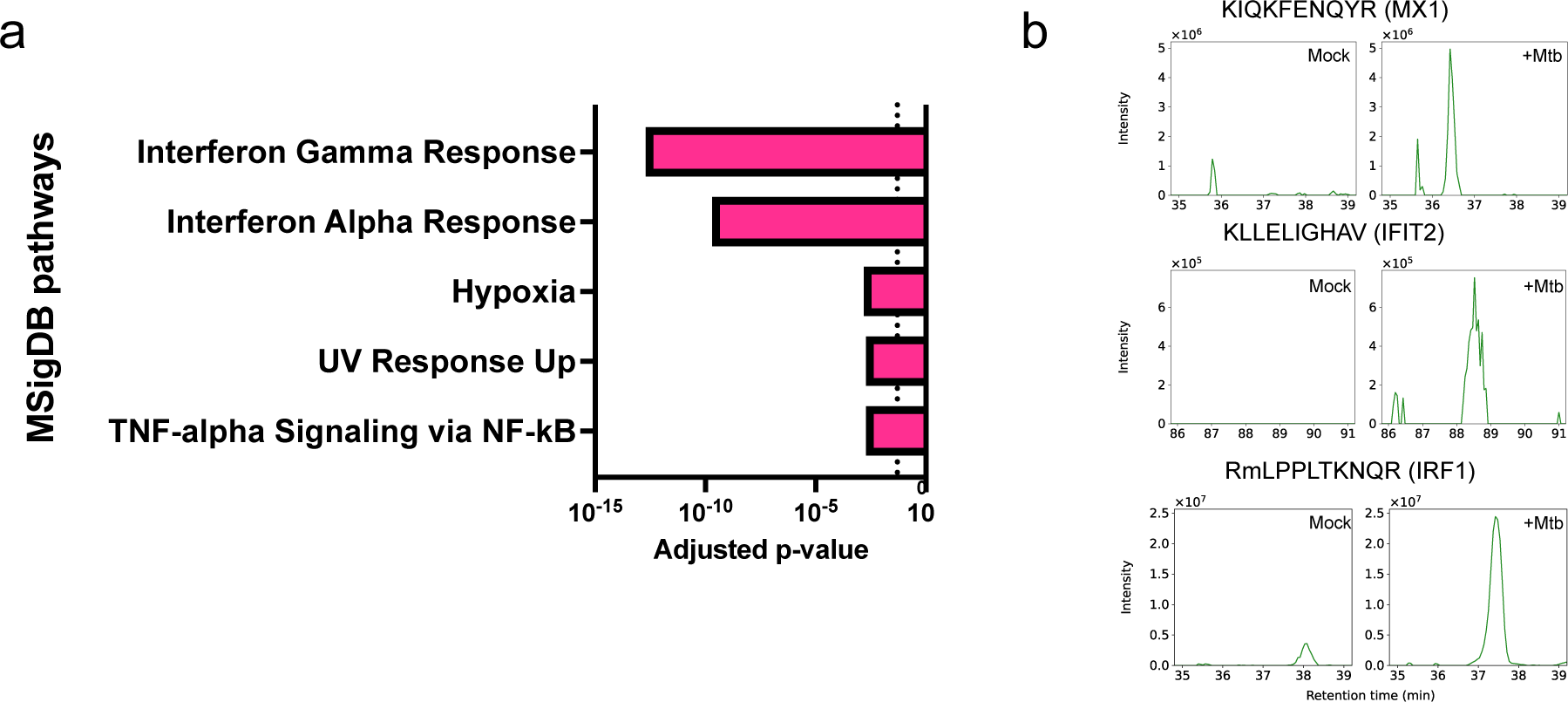
Self MHC-I peptides uniquely detected by PathMHC PRM analyses reflect the host cell response to *Mtb* infection. a) Top 5 enriched pathways among source proteins of self MHC-I peptides uniquely detected in PathMHC PRM analyses of *Mtb*-infected hMDCs from n = 3 donors (i.e., not identified in the initial DDA analysis). P-values were computed using Enrichr^50,51^ over the MSigDB Hallmark 2020 pathway database,^52^ using the set of source proteins of peptides detected in the corresponding DDA analyses of the *Mtb*-infected cells as a background. The dotted line marks an adjusted p-value threshold of 0.05. b) Representative extracted ion chromatograms for precursor ions of MHC-I peptides uniquely identified in PathMHC PRM analyses of *Mtb*-infected hMDCs.

## Discussion

PathMHC represents an approach to immunopeptidomics that is specifically tailored to addressing the challenges of vaccine development for infectious disease. PathMHC accelerates vaccine target discovery by computationally enriching for infection-specific precursor ions in discovery immunopeptidomics analyses, helping to find the “needle in a haystack” of pathogen-derived peptides in the MHC repertoire. In a model system where *Msmeg*-derived peptides make up ∼1% of the MHC-II immunopeptidome, PathMHC enhanced identification of *Msmeg-* derived peptides by 91% relative to a benchmark method based on DDA alone. The PathMHC computational workflow also generalizes to multiple pathogens, including SARS-CoV-2 infection and *Listeria monocytogenes* infection. Applying PathMHC to analyze the MHC-I repertoire of *Mtb*-infected hMDCs revealed that substrates of type VII secretion systems were selectively presented on MHC-I. Self peptides targeted by PathMHC reflected the host cell response to infection, particularly interferon responses.

PathMHC prioritizes pathogen-derived peptides for acquisition, enabling efficient use of instrument cycle time. By narrowing down the set of possible targets that must be scanned, PathMHC may also allow longer ion accumulation times for each peptide in settings where sensitivity is limiting. Further improvements to the computational pipeline will likely increase its ability to enrich for pathogen-derived peptides. DeepRTAlign and AutoMS were not originally trained on LC-MS data from immunopeptidomics studies, and retraining or fine-tuning the machine learning models underlying these tools on more task-specific data or substituting purpose-built tools may increase performance.

For some applications, PathMHC may have advantages over offline fractionation of MHC peptides as a means of increasing depth of coverage in immunopeptidomics studies. Offline fractionation can incur substantial sample losses,^39^ so PathMHC may have advantages when sample input is limited, such as when working with primary cells or tissues. PathMHC also requires less total instrument time, requiring 2 analyses per condition (4 hours) compared to 12 fractions^22^ (24 hours). Higher throughput could enable broader coverage of HLA alleles or pathogen variants, or could enable a faster response to emerging infectious disease risks.

Whereas here we benchmarked PathMHC against DDA as a baseline, library-free data-independent analysis (DIA) is also sometimes used as a method for untargeted discovery of MHC peptides.^40^ Unlike in DDA, all precursor ions are in principle fragmented and analyzed in DIA analyses, obviating the problem of prioritizing pathogen-derived precursors for MS/MS acquisition. However, DIA relies on accurate deconvolution of complex MS/MS spectra comprising fragment ions derived from many precursors. This can be a challenging computational task to perform reliably given the large search space of possible MHC peptides, unless constrained by prior knowledge in the form of a well-defined reference spectral library.^41^ A mismatch between the proportion of pathogen-derived peptides in the search space and the ground-truth proportion of pathogen-derived peptides in the sample can further increase false discovery rates for pathogen-derived peptides.^42^ Augmenting DIA-based discovery approaches with a workflow conceptually analogous to PathMHC in which candidate pathogen-specific peptides are secondarily screened by PRM could help confidently identify pathogen-derived peptides.

By applying PathMHC to identify *Mtb*-derived peptides presented on MHC-I, we showed that infected hMDCs preferentially present T7SS substrates on MHC-I and identified several antigens that could potentially serve as vaccine targets. Lewinsohn et al.^43^ previously isolated human CD8+ T cell clones from individuals with prior *Mtb* exposure that recognized 3 MHC-I epitopes derived from EsxB (AEMKTDAATL, EMKTDAATL, RADEEQQQAL) and 1 epitope conserved in both EsxG and EsxS (LLDAHIPQL) that we identified in this study. These overlapping results confirm that MHC-I antigens identified by MS can be immunogenic *in vivo* in humans.

CD8+ T cell responses against epitopes derived from non-T7SS antigens such as Ag85B have previously been reported, including epitopes that are restricted by HLA alleles expressed by cells used in this study.^44^ While we cannot rule out that these peptides went undetected in our study for technical reasons such as low ionization efficiency, the fact that we were unable to detect these antigens despite using PathMHC to increase coverage of infection-specific peptides suggests that presentation of T7SS substrates is favored in hMDCs for biological reasons and the low frequency of *Mtb*-derived MHC-I peptides reflects real bottlenecks in access to antigen processing pathways. It is possible that CD8+ T cell responses against other classes of antigens could be primed by antigen presenting cell types other than monocyte-derived phagocytes, or by cells that have taken up dead *Mtb* material rather than live *Mtb*. If this were the case, CD8+ T cells targeting T7SS substrates might be more effective at recognizing the monocyte-derived cells that make up a large proportion of infected cells in TB granulomas.^34^

*Mtb* infects a range of phagocytic human immune cells, including both macrophages and dendritic cells.^34,35^ Hypothetically, these cell types might express and use different antigen processing pathways and therefore present different sets of *Mtb* antigens on MHC-I. Heterogeneous antigen presentation would pose an obstacle to efficient clearance of the infection by CD8+ T cells, as T cells capable of recognizing one cell type might be unable to recognize the other. However, we previously showed that T7SS substrates predominate among *Mtb*-derived MHC-I antigens presented by human monocyte-derived macrophages,^22^ and our results here show that the same is true of hMDCs. Some MHC-I peptides (LLDEGKQSL from EsxA and QEQASQQIL from EsxJKPW) and source proteins (EsxB, EspC) were identified in both studies. T cells primed to recognize T7SS substrates may be able to recognize both *Mtb*-infected monocyte-derived dendritic cells and macrophages, making T7SS substrates a promising class of potential vaccine targets.

The workflow underlying PathMHC can in principle be extended to immunopeptidomics in non-infectious disease areas, and to other types of mixed host-pathogen proteomic samples. For example, a workflow analogous to PathMHC could potentially be used to identify MHC peptides uniquely presented following treatment of tumor cells with a therapy to nominate potential targets for combination immunotherapy,^45,46^ or could be used to specifically identify pathogen-derived interaction partners of a host protein of interest in IP-MS or AP-MS studies to better understand host-pathogen interactions.^47^

In addition to pathogen-derived peptides, PathMHC enriches for infection-associated MHC peptides derived from self proteins upregulated during the host cell response to infection. While in general these infection-associated peptide-MHC complexes could in principle be targets for the design of therapies and diagnostics, targeting the interferon-associated MHC peptides detected by PathMHC in *Mtb*-infected cells could result in low specificity and off-target effects since these peptides could be presented by any cell responding to interferon. Targeting these peptides using vaccines would likely be difficult, as dedicated mechanisms in the thymus ensure deletion of T cells specific for infection-induced self MHC peptides.^48^ Using uninfected control samples treated with stimuli that mimic the host response to infection (such as recombinant type I interferon in the case of *Mtb*) could help ensure that a higher proportion of differentially presented antigens targeted by PathMHC are truly pathogen-specific peptides rather than infection-induced self peptides. In other types of host-pathogen proteomic samples besides MHC peptides, infection-associated host proteins may be of greater biological interest and could reveal novel information about the host response to infection.

Our results show that computational enrichment of infection-specific precursor ions enables immunopeptidomics studies to take advantage of the specificity and sensitivity of targeted MS to more efficiently detect pathogen-derived peptides without prior knowledge of the identities of target epitopes. Selectively targeting infection-associated peptides in discovery immunopeptidomics studies can potentially make target discovery for T cell priming vaccines more efficient across a range of infectious disease areas.

## Materials and methods

### M. tuberculosis *culture*

*Mycobacterium tuberculosis* (*Mtb*) H37Rv was grown in Difco Middlebrook 7H9 media supplemented with 10% OADC, 0.2% glycerol, and 0.05% Tween-80 to mid-log phase.

### M. smegmatis *culture*

*Mycobacterium smegmatis* (*Msmeg*) mc^2^155 was grown in Difco Middlebrook 7H9 media supplemented with 10% OADC, 0.2% glycerol, and 0.05% Tween-80 overnight (12-14 hours).

### Human cell isolation, differentiation, and culture

Deidentified buffy coats were obtained from Massachusetts General Hospital, except for donor C (provided by StemCell). Samples are acquired and provided to research groups with no identifying information. PBMCs were isolated by density-based centrifugation using Ficoll (GE Healthcare). CD14 +monocytes were isolated from PBMCs using a CD14 positive-selection kit (Stemcell). Isolated monocytes were differentiated in R10 media [RPMI 1640 without phenol red (Gibco) supplemented with 10% heat-inactivated FBS (Gibco), 1% HEPES (Corning), 1% L-glutamine (Sigma)] supplemented with 25 ng/mL GM-CSF (Biolegend, 572902) and 25 ng/mL IL-4 (Biolegend, 574006). Monocytes were cultured on tissue culture treated T75 flasks (VWR) for 6 days. Media was replaced with fresh cytokine-containing media every 3 days.

### HLA genotyping

Genomic DNA was extracted from 5×10^6^ PBMCs using a Qiagen DNeasy kit. HLA typing was performed using a targeted next generation sequencing (NGS) method. Briefly, locus-specific primers were used to amplify a total of 26 polymorphic exons of HLA-A & B (exons 1–4), C (exons 1–5), E (exon 3), DPA1 (exon 2), DPB1 (exons 2–4), DQA1 (exon 1–3), DQB1 (exons 2 & 3), DRB1 (exons 2 & 3), and DRB3/4/5 (exon 2) genes with Standard BioTools Access Array system (Standard BioTools, South San Francisco, CA 94080 USA). The 26 Standard BioTools PCR amplicons were harvested from Standard BioTools Access Allay IFC and pooled. Quality and quantity were checked using a Caliper LabChip GX Touch HT Nucleic Acid Analyzer (PerkinElmer, Waltham, MA 02452 USA). The PCR product library was quantitated and subjected to sequencing on an Illumina MiSeq sequencer (Illumina, San Diego, CA 92122 USA). HLA alleles and genotypes were called using the Omixon HLA Explore (version 2.0.0) software (Omixon Biocomputing Ltd., Budapest, Hungary). HLA genotyping data for donor C was instead provided by StemCell.

### M. smegmatis infection

The *Msmeg* culture was pelleted by centrifugation, washed once with PBS, and resuspended in R10 media. 25 million hMDCs were infected at a multiplicity of infection (MOI) of 10 for 2 hr (or mock-infected with media not containing *Msmeg*) and then washed with PBS to remove extracellular *Msmeg*. Infected hMDCs were cultured in R10 media for 24 hr before harvesting.

### M. tuberculosis infection

The *Mtb* culture was pelleted by centrifugation, washed once with PBS, resuspended in R10 media and centrifuged at low speed (500 rpm for 5 minutes) to pellet clumps, leaving a uniform suspension of bacteria in the supernatant. 50 million hMDCs were infected at MOI 2.5 for 4 hr and then washed with PBS to remove extracellular *Mtb*. Infected hMDCs were cultured in R10 media for 72 hours before harvesting.

### MHC immunoprecipitation

hMDCs were harvested by collecting the culture media (containing any cells in suspension), washing adherent cells with PBS, incubating remaining adherent cells with PBS supplemented with 4 mM EDTA for 15 minutes at 37 °C, gently scraping with a cell scraper, collecting the detached cells, and washing the flask with PBS and collecting the wash. The harvested cells were then pelleted, washed with PBS, and lysed in 1 mL of MHC lysis buffer [20 mM Tris, 150 mM sodium chloride, pH 8.0, supplemented with 1% CHAPS, 1 x HALT protease and phosphatase inhibitor cocktail (Pierce), and 0.2 mM phenylmethylsulfonyl fluoride (Sigma-Aldrich)].

#### BSL3 protocol

Lysate from *Mtb*-infected (or mock-infected) cells was sonicated using a Q500 ultrasonic water bath sonicator (Qsonica) in five 30 second pulses at an amplitude of 60%, cleared by centrifugation at 16,000 x g for 5 minutes, and sterile filtered twice using 0.2 μm filter cartridges (Pall NanoSep).

#### BSL2 protocol

Lysate from *Msmeg*-infected (or mock-infected) cells was sonicated using a VCX-130 probe ultrasonic processor (Sonics) in three 10 second pulses at an amplitude of 30% and cleared by centrifugation at 16,000 x g for 5 minutes.

Lysates were then added to protein A sepharose beads pre-conjugated with 0.5 mg of pan-MHC-I antibody (clone W6/32) or 0.1 mg of pan-MHC-II antibody (clone Tü39), prepared as previously described,^46^ and incubated rotating at 4 °C overnight (12–14 hours). Beads were then washed and peptide-MHC complexes eluted as previously described.^46^

### MHC-I peptide isolation

C18 SpinTips (Protea) were washed with 0.1% trifluoroacetic acid, activated with 90% acetonitrile supplemented with 0.1% formic acid, and washed with 0.1% formic acid. Eluate from MHC-I IPs was applied to the column by centrifugation. The column was washed with 0.1% formic acid, and peptides were eluted by applying elution solvent (28% acetonitrile with 0.1% formic acid) by centrifugation twice. Eluates were snap-frozen in liquid nitrogen and lyophilized.

### MHC-II peptide isolation

MHC-II-associated peptides were purified using 10 kDa molecular weight cutoff filters (Cytiva NanoSep) as previously described,^46^ snap-frozen in liquid nitrogen, and lyophilized.

### MS analyses

For all MS analyses, samples were analyzed using an Orbitrap Exploris 480 mass spectrometer (Thermo Fisher Scientific) coupled with an UltiMate 3000 RSLC Nano LC system (Dionex), Nanospray Flex ion source (Thermo Fisher Scientific), and column oven heater (Sonation). The MHC peptide sample was loaded onto a fused silica capillary chromatography column with an integrated electrospray tip (∼1 μm orifice) prepared and packed in-house with 10 cm of 1.9 μm C18 beads (ReproSil-Pur). MHC peptide samples were resuspended in 0.1% formic acid and loaded at a flow rate of 300 nL/minute. Peptides were eluted using a flow rate of 100 nL/minute.

Standard mass spectrometry parameters were as follows: spray voltage, 2.0 kV; no sheath or auxiliary gas flow; ion transfer tube temperature, 275 °C. MHC-I peptides were eluted using a gradient of 6–25% buffer B (70% Acetonitrile, 0.1% formic acid) over 75 min, 25–45% over 5 min, 45–100% over 5 min, hold for 1 min, and 100% to 3% over 2 min. MHC-II peptides were eluted using a gradient of 6–12% buffer B (70% Acetonitrile, 0.1% formic acid) over 10 min, 12– 20% over 45 min, 20–25% over 10 min, 25-45% over 5 minutes, 45-97% over 2 minutes, hold for 1 min, and 97% to 3% over 2 min. Full scan mass spectra (120,000 resolution 350–1200 m/z for MHC-I, 350-1800 m/z for MHC-II) were detected in the orbitrap analyzer after accumulation of 3×10^6^ ions (normalized AGC target of 300%) or 25 ms. For every full scan, MS/MS scans were collected during a 3 s cycle time. Ions were isolated (0.4 m/z isolation width) using the standard AGC target and automatic determination of maximum injection time, fragmented by HCD with 30% CE, and scanned at a resolution of 120,000. Charge states <2 and >4 were excluded. In DDA analyses, precursors were excluded from selection for 30 seconds if fragmented n=2 times within a 20-second window.

For PathMHC PRM analyses, inclusion list masses were matched by m/z and charge state, with a mass tolerance of 5 ppm. Acquisition was scheduled for a window of 3 minutes centered on the observed retention time of the target precursor ion in the DDA analysis in which it was originally detected. Precursors were excluded from selection for 4 seconds after being fragmented 1 time.

For analyses of MHC-II peptides from hMDCs infected with *Msmeg*, 50% of a given sample was injected for each DDA analysis and 50% for a follow-up PRM or DDA analysis. For analyses of MHC-I peptides from hMDCs infected with *Mtb*, 40% of the sample was used for each analysis while 20% was reserved for subsequent validation using SureQuant.

DDA-only MHC-I analyses of MHC-I peptides from *Mtb*-infected hMDCs were performed without prior offline fractionation (donor E), or with offline fractionation by reversed-phase HPLC as previously described^22^ (donor D).

### PathMHC computational pipeline

The PathMHC pipeline first identifies MS1 peaks (features) using Dinosaur (version 1.2.0), then aligns these features between a sample from infected cells and a mock-infected control using DeepRTalign (version 1.1.3). MS1 features detected in the infected sample are excluded if they are mapped to a feature in the mock-infected control by DeepRTalign, or if they match a peptide already identified by DDA from a provided list of peptide-spectrum matches to within a mass tolerance of 10 ppm and retention time tolerance defined by the peak boundaries set by Dinosaur. The remaining MS1 features (i.e., those not previously identified by DDA and with no corresponding peak in the mock-infected control) are then scored using AutoMS, using an XIC length of 40 seconds and an XIC mass tolerance of 10 ppm for MHC-I peptides and 40 ppm for MHC-II peptides. A final inclusion list is generated consisting of precursors meeting the following criteria: For MHC-I, charge state 2-4, retention time less than 90 minutes (from the end of sample loading), AutoMS signal to noise ratio score (SNR) > 1 or AutoMS score > 0.2. For MHC-II, charge state 2-3, retention time less than 90 minutes, AutoMS signal to noise ratio (SNR) > 1.5 or AutoMS score > 0.3.

### Proteome Discoverer PathMHC computational pipeline

To implement an analogue of our PathMHC computational pipeline in Proteome Discoverer, we created a workflow in which LC-MS features are detected using Minora in the processing step and aligned between the infected and mock-infected sample using the Feature Mapper (Perform RT Alignment = True, Max RT Shift = 10 min, Min S/N Threshold = 2) and Precursor Ions Quantifier nodes in the consensus step. Consensus features are then exported with the following filters applied: charge state = 2 or 3, intensity in mock-infected control sample = 0, average apex retention time is between 25 minutes and 115 minutes (should be set depending on your LC gradient and duration of sample loading). The resulting table of precursor ions is reformatted for use as an inclusion list for PRM (see https://github.com/oleddy/pathMS).

### MS data search and manual inspection

All mass spectra were analyzed with Proteome Discoverer (PD, version 3.0) and searched using Sequest with rescoring using INFERYS and Percolator against a custom database comprising the Uniprot human proteome (UP000005640) together with either the Uniprot *Mycobacterium tuberculosis* H37Rv proteome (UP000001584) or the *Mycobacterium smegmatis* mc^2^155 proteome (UP000006158). No enzyme was used, and variable modifications included oxidized methionine for all analyses. Peptide-spectrum matches from MHC-I analyses were filtered with the following criteria: search engine rank = 1, length between 8 and 13 amino acids, XCorr ≥ 2.0, spectral angle ≥ 0.6, and percolator q-value < 0.05. Peptide-spectrum matches from MHC-II analyses were filtered with the following criteria: search engine rank = 1, length between 8 and 30 amino acids, XCorr ≥ 2.0, spectral angle ≥ 0.6, and percolator q-value < 0.05.

Identifications (IDs) of putative *Mtb*-derived peptides were rejected if any peptide-spectrum matches (PSMs) for the same peptide were found in the unfiltered DDA MS data for the corresponding mock-infected control. For each putative Mtb peptide identified, MS/MS spectra and extracted ion chromatograms (XIC) were manually inspected, and the ID was only accepted for further validation if it met the following criteria: (1) MS/MS spectra contained enough information to unambiguously assign a majority of the peptide sequence; (2) neutral losses were consistent with the chemical properties of the peptide; (3) manual de novo sequencing did not reveal an alternate peptide sequence that would explain a greater number of MS/MS spectrum peaks; (4) XIC showed a peak in MS intensity at the mass to charge ratio (m/z) of the peptide precursor ion at the retention time at which it was identified that did not appear in the corresponding mock-infected control. Peptides that met these criteria were further validated using SureQuant (see below).

### Synthetic standard survey MS analyses

DDA MS analysis of the SIL peptide mixture was performed as described above (see MS analyses) with the following modifications: Peptides were eluted using a gradient of 6–35% buffer B over 30 min, 35–45% over 2 min, 45–100% over 3 min, and 100% to 2% over 1 min. No dynamic exclusion was used.

A second set of survey analyses was performed on the mixture of SIL peptides with background matrix using the full SureQuant acquisition method (see below). SIL peptides were spiked into a mixture of MHC-I peptides purified as described above from THP-1 cells differentiated into macrophages via 24 hr of treatment with 150 nM phorbol myristate acetate (PMA), which provided a representative background matrix. Because SIL amino acids are not 100% pure, SIL peptide concentrations were adjusted and survey analyses were repeated until the SIL peptide could be reliably detected while minimizing background signal detected at the mass of the biological peptide. In the final SIL standard mixture, 100 fmol of each peptide was used per sample for all epitopes except two peptides for which 10 fmol was used per sample (KIYSEADEAWRK and RADEEQQQAL).

### Validation of candidate Mtb peptides by SureQuant MS analysis

Stable isotope labeled (SIL) synthetic peptide standards were ordered from BioSynth as a crude peptide library. Standard MS parameters and MS1 scan parameters were as described above, except that a scan range of 380-1200 m/z was used. The custom SureQuant method was built based on the template provided in the Thermo Orbitrap Exploris Series 2.0 method editor. After the optimal charge state and most intense product ions were determined via a survey analysis of the synthetic SIL peptide standards alone (see above), a method branch was created for each m/z offset between the SIL peptide and biological peptide as previously described.^49^ The m/z tolerance for detection of SIL standards was set at 10 ppm. A resolution of 15,000 and automatically determined ion accumulation time was used for MS/MS scans of the SIL standards, and a resolution of 240,000 with a maximum accumulation time of 1 s was used for MS/MS scans of the biological peptide. An MS/MS scan of the light (biological) peptide was triggered upon detection of 3 or more of the top 6 product ions (mass tolerance 20 ppm) in a scan of the corresponding SIL standard. Results were analyzed using Skyline Daily Build 24.1.1.202.

### Analysis of publicly available immunopeptidomics datasets

Raw LC-MS/MS data underlying MHC-I immunopeptidomics datasets from *Listeria*-infected cells and SARS-CoV-2-infected cells generated by Mayer et al.^20^ and Weingarten-Gabbay et al.^21^ respectively were obtained from their respective submissions to the ProteomeXchange PRIDE database. In the case of Mayer et al.’s dataset, for each biological replicate raw MS data from *Listeria*-infected and uninfected cells was analyzed using the PathMHC computational pipeline as described above, with the exception that precursor ions with existing peptide-spectrum matches (PSMs) were not excluded. Putatively infection-specific precursor ion peaks selected by the pipeline were matched with reported PSMs with an m/z tolerance of 10 ppm. The proportion of putative *Listeria-*dervied MHC-I peptides among identified precursors selected by the pipeline was compared with the total proportion of *Listeria*-derived peptides in the dataset. One of four provided replicates was excluded because one of the files provided could not be analyzed by Dinosaur, possibly due to an incompatible file format or corruption of the file. The data from Weingarten-Gabbay et al. was analyzed similarly, but in this dataset each replicate represented paired peptide fractions from offline fractionation rather than biological replicates. For this dataset, the proportion of SARS-CoV-2-derived peptides enriched by the PathMHC computational pipeline was compared to the overall proportion of SARS-CoV-2 peptides for each replicate as PSMs were reported separately for each fraction. In addition to a 10 ppm m/z tolerance, a 1.5-minute retention time tolerance was used, as retention times for each PSM were also reported.

### Gene set enrichment analysis

Self MHC-I peptides uniquely identified in PathMHC PRM analyses of *Mtb*-infected hMDCs that had not been identified by DDA were mapped to their source proteins. This set of proteins was analyzed using Enrichr,^50,51^ using the set of source proteins of self MHC-I peptides identified by DDA as a background. The MSigDB Hallmark 2020 pathway database was used for pathway enrichment analysis.^52^ Pathway enrichment analyses of self peptides based on computational analyses of publicly available immunopeptidomics datasets using the PathMHC computational pipeline were performed analogously. The precursors selected by PathMHC that had associated PSMs were mapped to their source proteins, and these proteins were analyzed using Enrichr as described above, using the overall set of proteins identified in the analysis as a background.

### Protein secretion pathway enrichment analysis

To determine whether *Mtb*-derived MHC-I peptides or their source proteins were enriched for proteins with specific secretion signals, we classified each protein in the reference *Mtb* H37Rv proteome (UP000001584) using SignalP 6.0,^53^ and added an additional class of known T7SS substrates (all Esx-family proteins, all PE/PPE proteins, and EspA, EspB, EspC, EspE, EspF, EspI, EspJ, and EspK). Under a null model in which all possible *Mtb* peptides are equally likely to be presented, the probability of drawing a peptide from a given class of proteins is approximately the sum of the lengths of the amino acid sequences of proteins in that class divided by the total number of amino acid residues in the *Mtb* proteome. Using this approximation, we used a binomial test to assess whether the number of *Mtb*-derived peptides from each class of proteins detected in the MHC-I was greater or less than would be predicted under the null model. We used a hypergeometric test to assess whether the number of source proteins from each class was greater or less than expected, relative to a null model in which each protein in the *Mtb* proteome is equally likely to contribute to the MHC-I repertoire.

## Code availability

Our computational pipeline for PathMHC and all other code used in this study is available at https://github.com/oleddy/pathMS.

## Data availability

Raw mass spectrometry data have been deposited to the ProteomeXchange Consortium via the PRIDE with the dataset identifier PXD055726.^54^

## Acknowledgements

The authors thank all the staff members of the Ragon Institute, the Koch Institute, and MIT for the essential work they do to make our research possible. The Tü39 hybridoma used for in-house antibody production was generously provided by Claudia Falkenburger and Hans-Georg Rammensee. We thank Lauren Stopfer, Cameron Flower, Tigist Tamir, Elizabeth Choe, and Ryuhjin Ahn for helpful conversations, training, and technical guidance. We modified code written by Cameron Flower to plot MS/MS spectra. We thank the developers of Dinosaur, DeepRTalign, and AutoMS for making their software free and open-source, as well as the developers of other Python and R libraries used in this study. This work is supported by funding from the MIT Center for Precision Cancer Research at the Koch Institute and U.S. NIH grants U01 CA238720, U54 CA283114, 1R35GM142900, and R01A1022553. O.L. is supported by U.S. NIEHS grant T32-ES007020. This work was performed in part in the Ragon Institute BSL3 core facility, which is supported by the NIH-funded Harvard University Center for AIDS Research (P30 AI060354). We thank Yong Xie, Julie Boucau, and Eliane Shwairi for managing the facility.

---

**Supplementary Figure 1.**
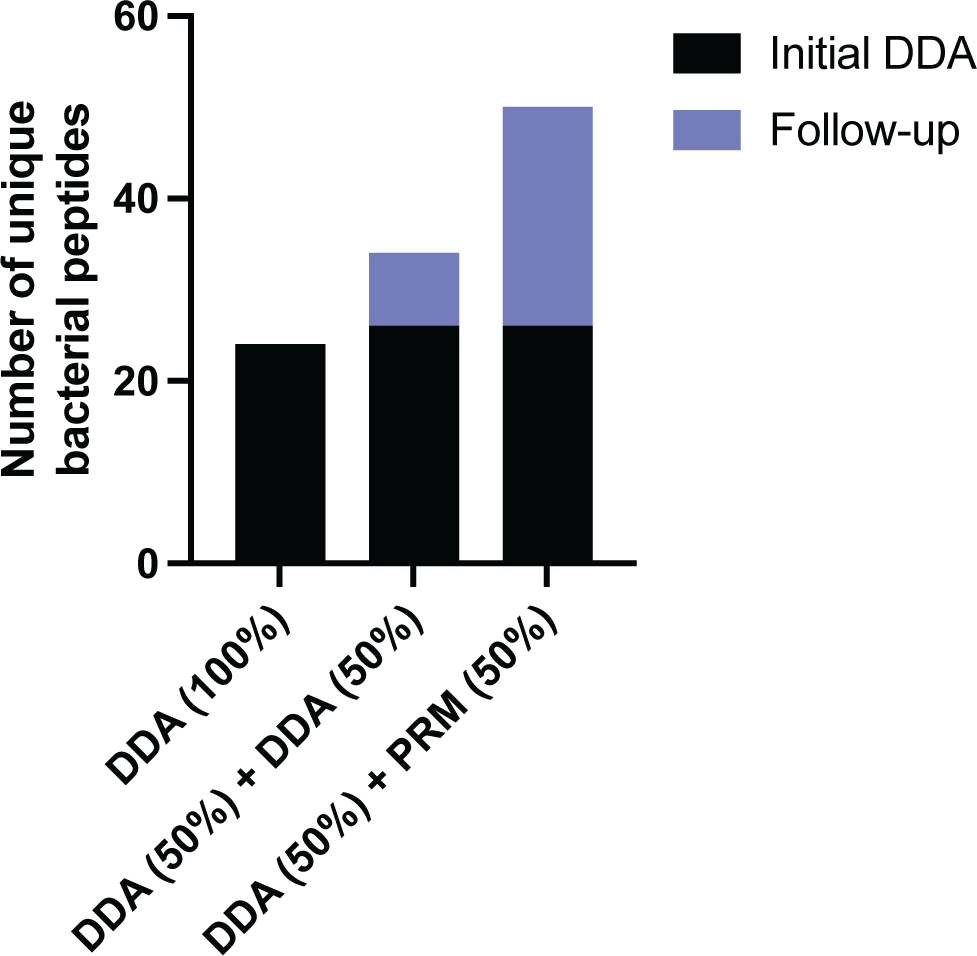
Identifying an appropriate benchmark for PathMHC analyses. Number of unique bacterial peptides identified in analyses of MHC-II peptides isolated from hMDCs infected with *Msmeg* using three workflows: 100% of sample injected in a single DDA analysis, 50% of sample injected in each of two DDA analyses, or 50% of sample injected in a DDA analysis and 50% in a PRM analysis using a prototype version of the PathMHC workflow.

**Supplementary Figure 2.**
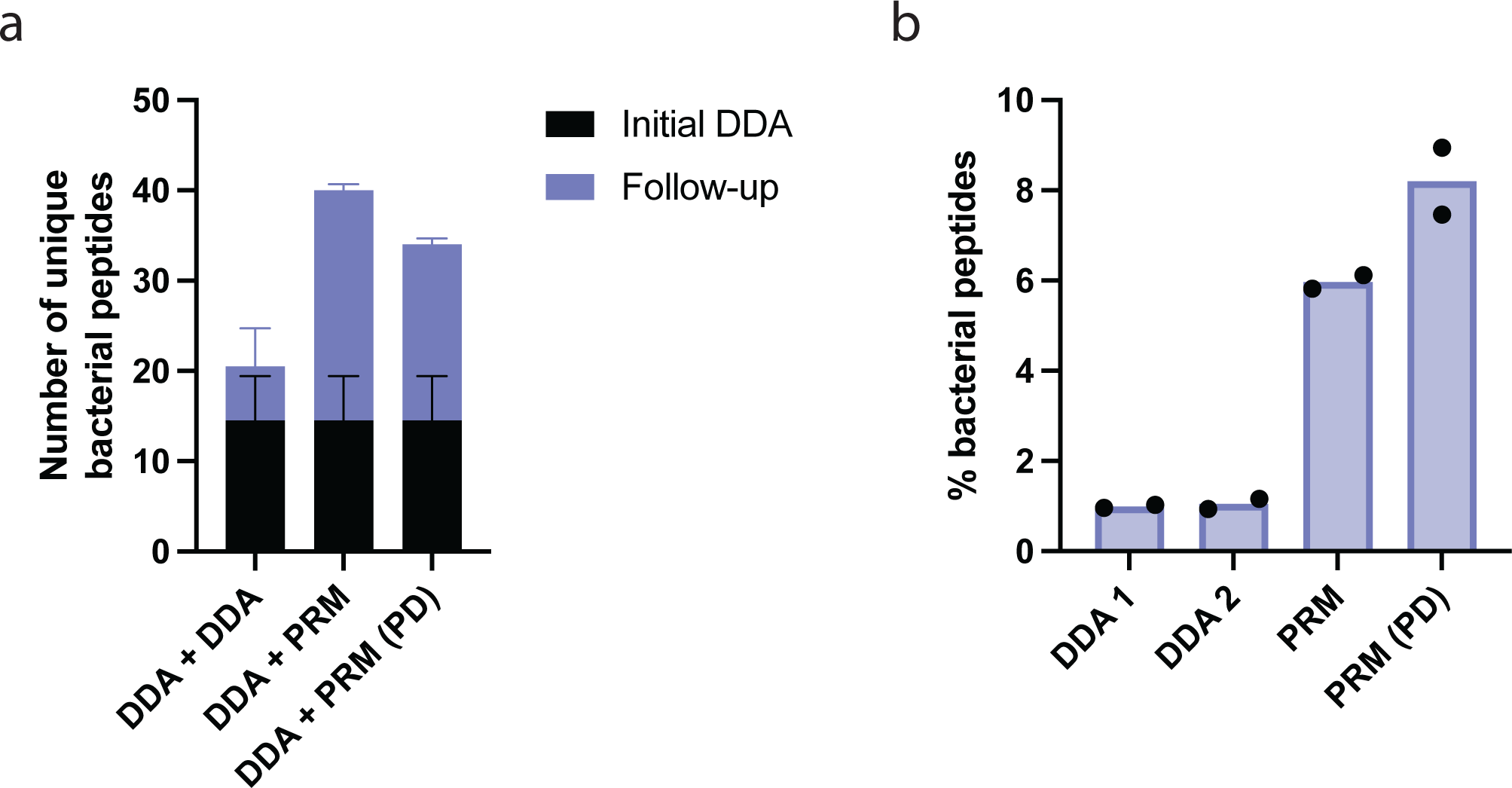
A PathMHC computational pipeline implemented in Proteome Discoverer and an open source implementation achieve comparable performance. Number of unique bacterial peptides (a) and percentage of bacterial peptides as a proportion of total peptides (b) identified in analyses of MHC-II peptides isolated from hMDCs from n = 2 donors infected with *Msmeg*, using a DDA-only benchmark protocol, a version of PathMHC using open-source software tools, and a version of PathMHC using a computational pipeline created in Proteome Discoverer (PD).

**Supplementary Figure 3.**
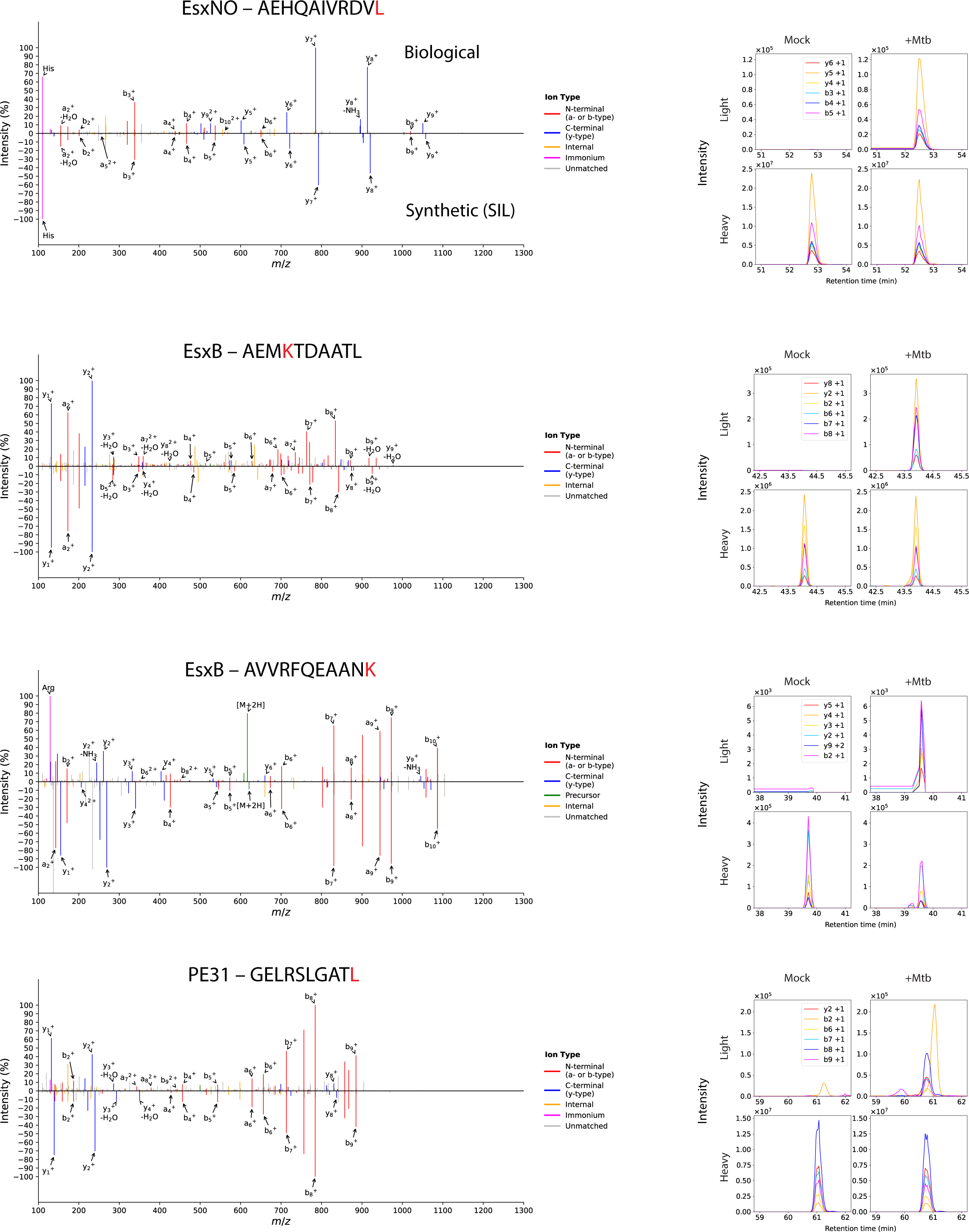
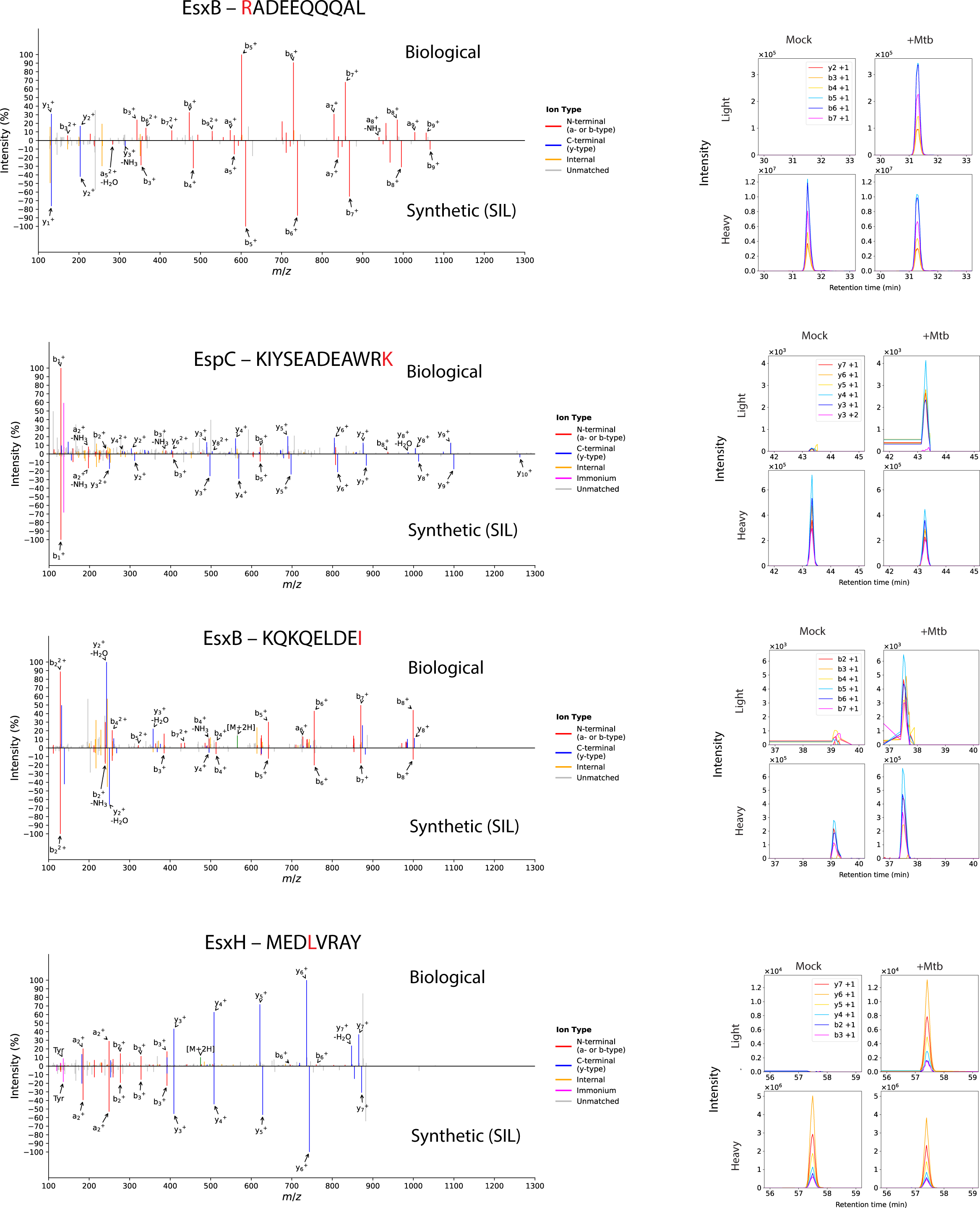
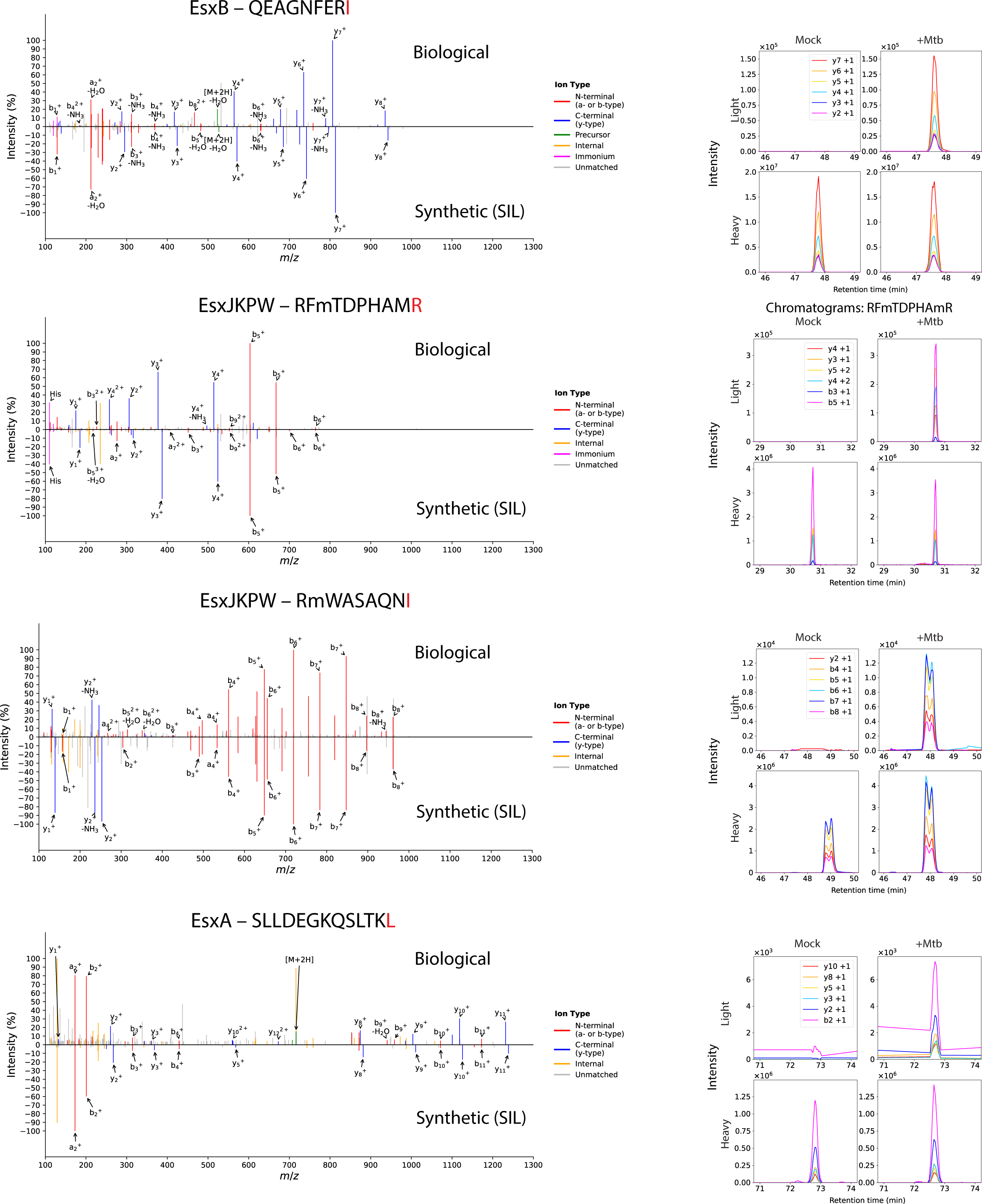
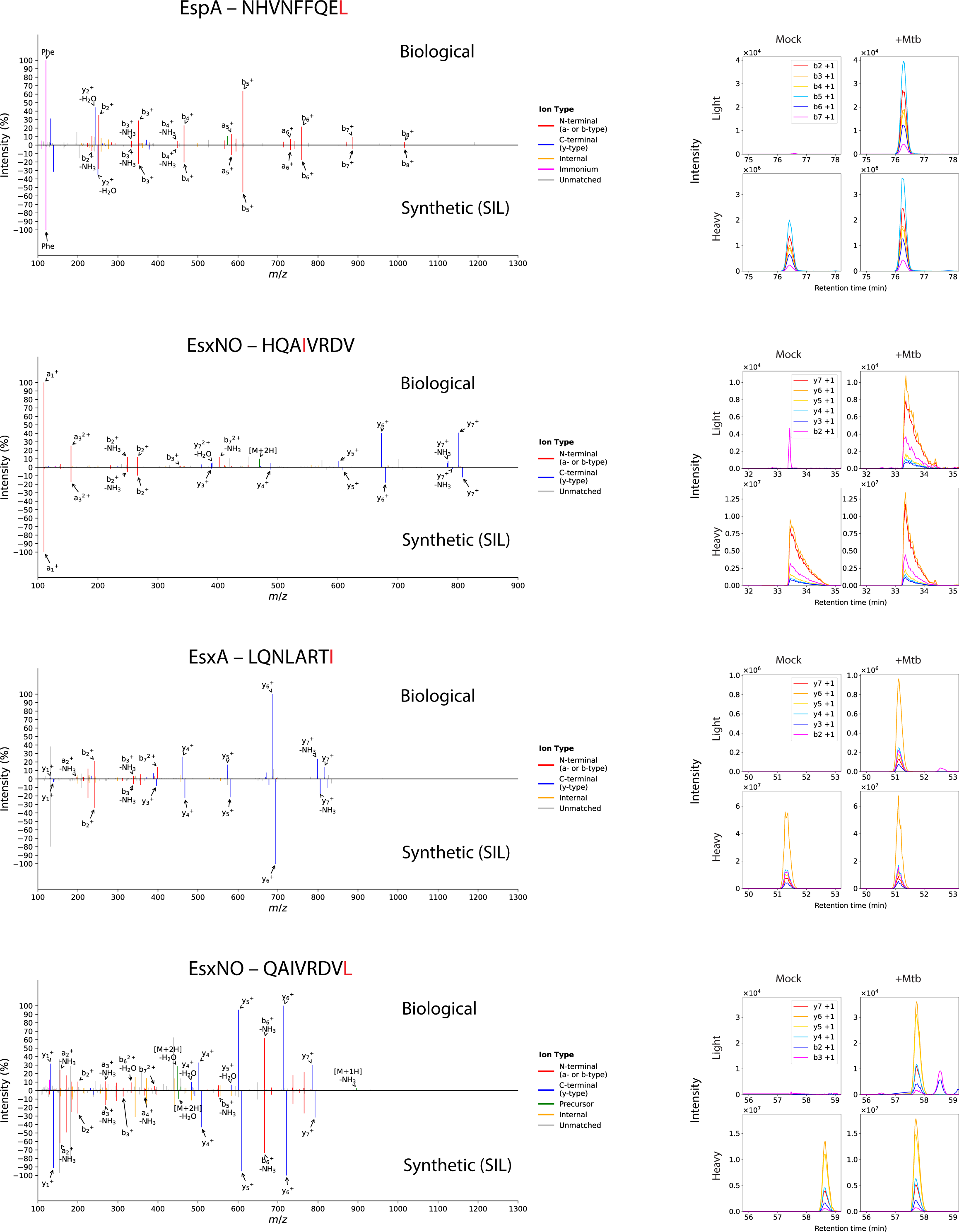
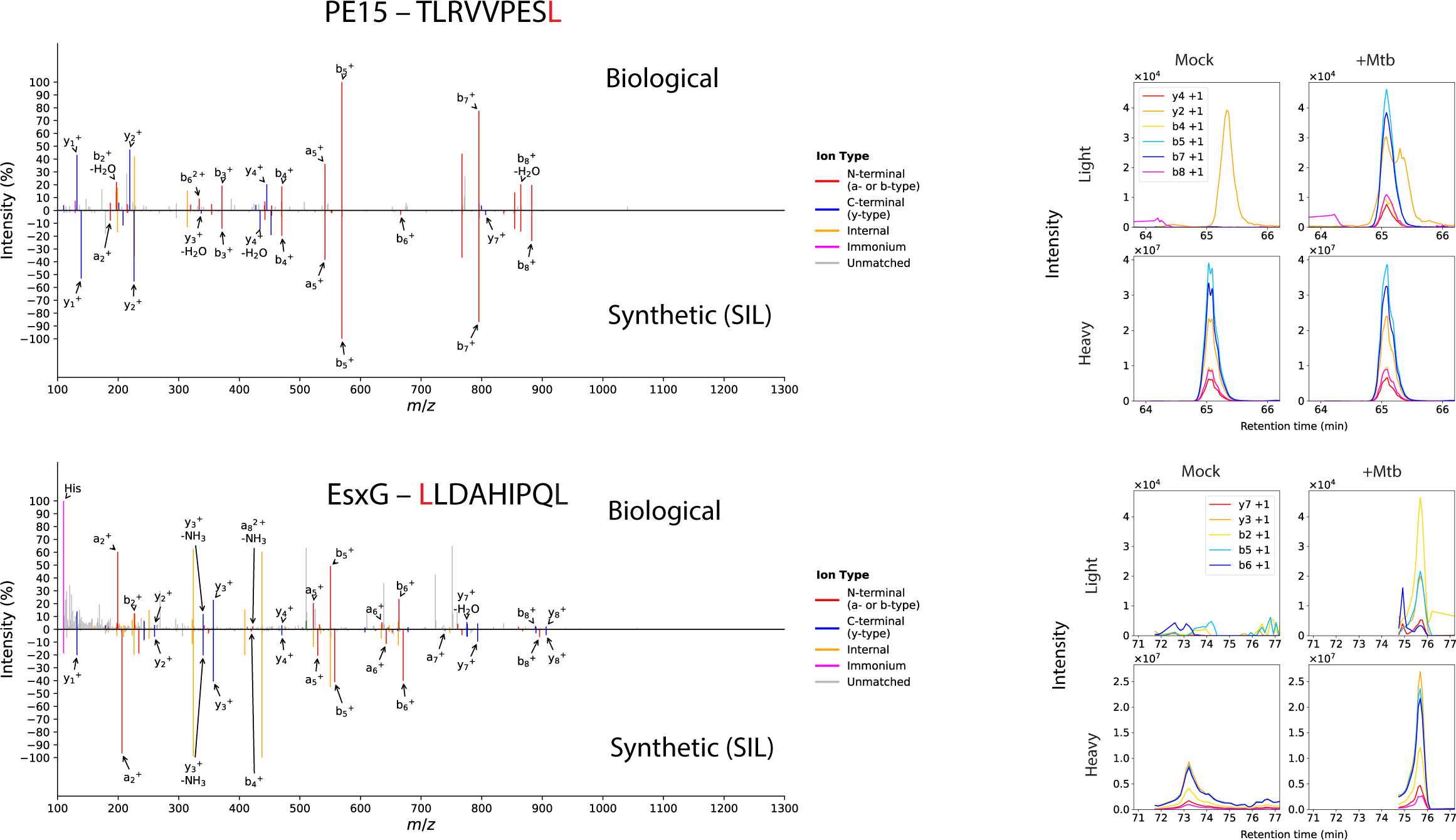
Validation of *Mtb*-derived MHC-I peptide identifications using SureQuant. MS/MS spectrum comparisons (left) and SureQuant product ion chromatograms for *Mtb*-infected and mock-infected hMDCs (right column) for each peptide. For each peptide, the amino acid that is stable isotope labeled in the synthetic standard is indicated in red.

**Supplementary Figure 4.**
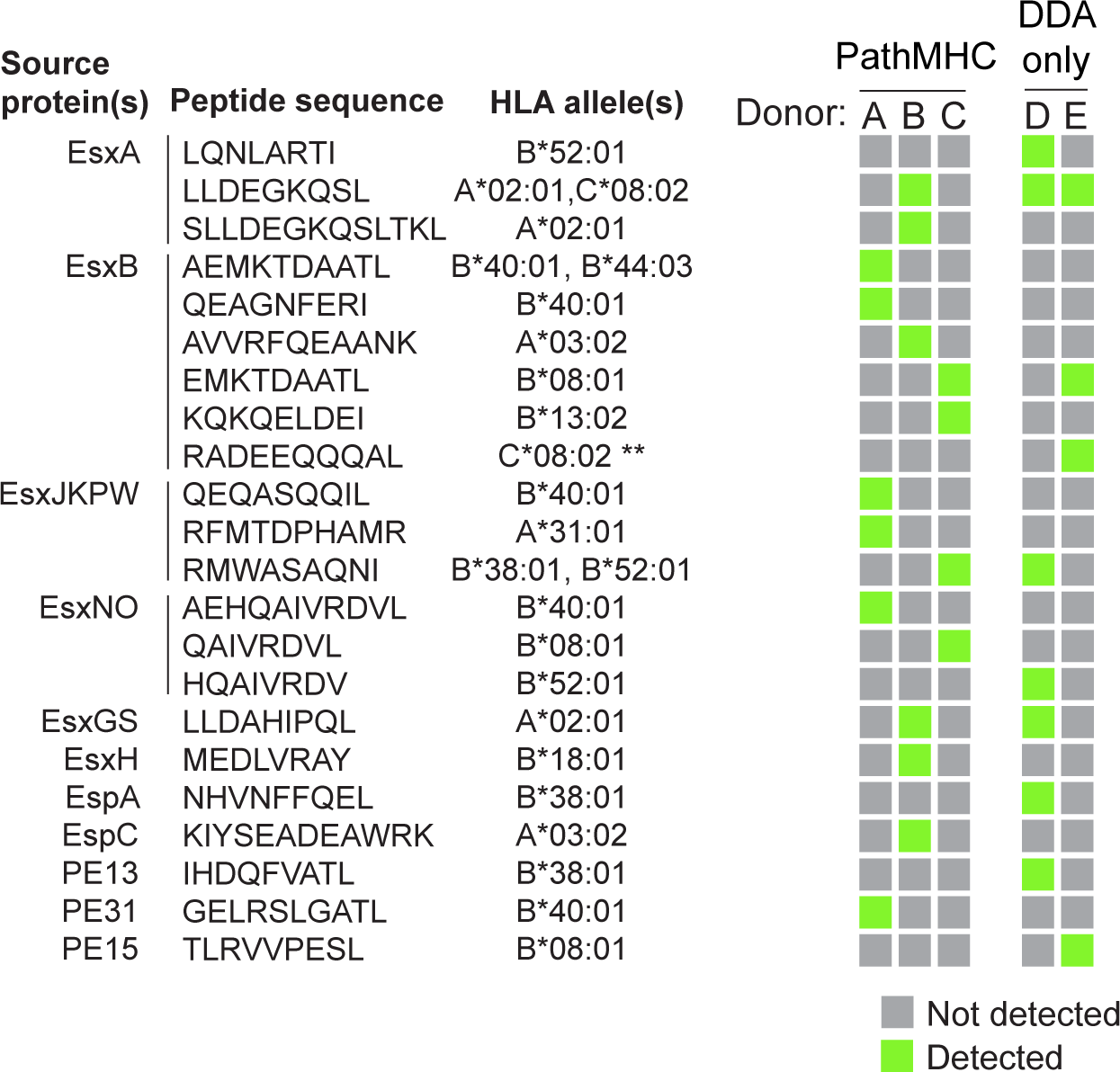
Summary of all *Mtb*-derived MHC-I peptides identified in infected hMDCs, including both PathMHC and DDA-only analyses. Sequences, source proteins, associated HLA alleles, and donors for each validated *Mtb*-derived MHC-I peptide. Green boxes indicate the donor(s) that presented a given peptide. The associated HLA alleles listed are the allele with the top Eluted Ligand (EL) rank predicted by NetMHCpan 4.1,^30^ for each donor that presented a given peptide. (** This epitope has previously been reported as an HLA-B*14-restricted CD8+ T cell epitope.^43^ HLA-B*14:02, expressed by donor E, may in fact be the restricting allele, but the peptide is not predicted to bind HLA-B*14:02 by NetMHCpan.)

**Supplementary Figure 5.**
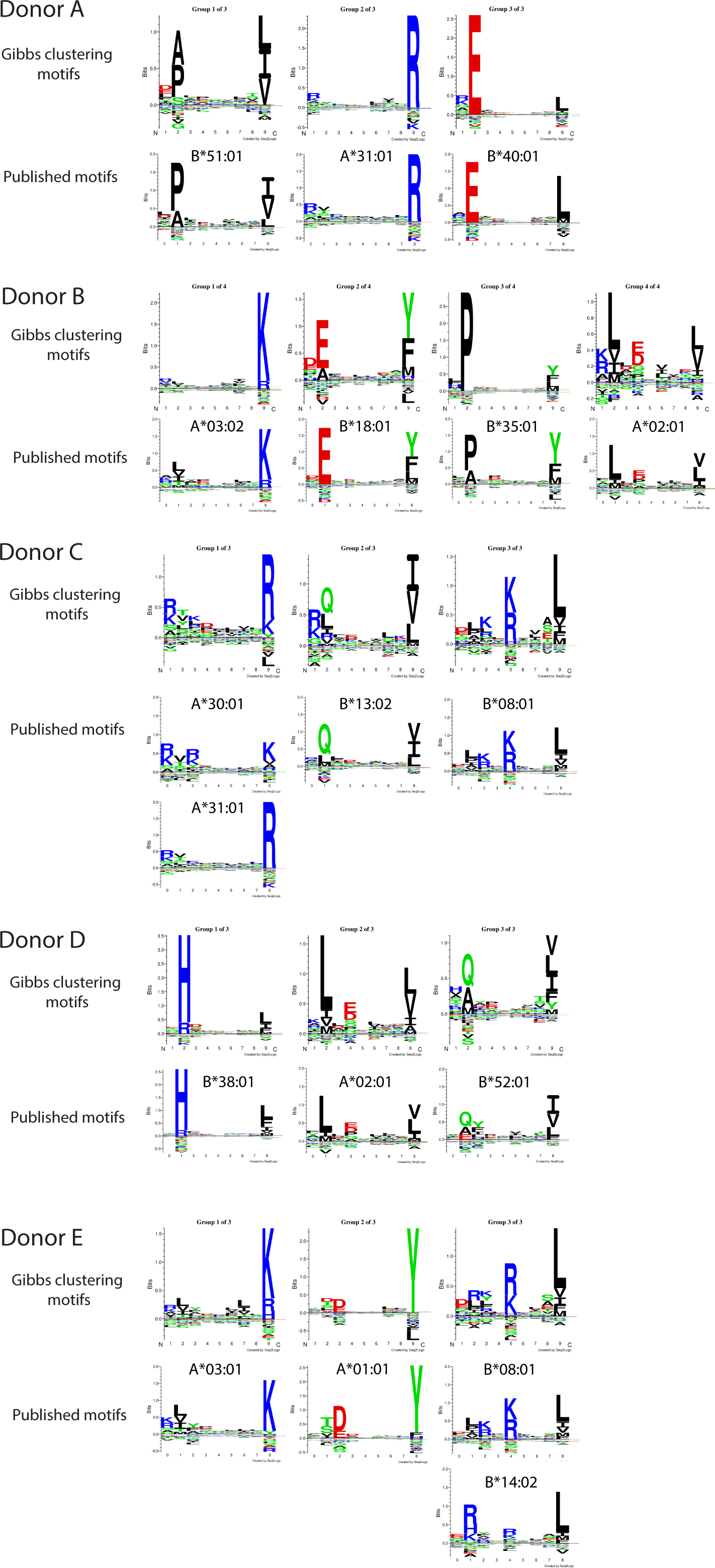
Gibbs clustering of MHC-I peptides presented by *Mtb*-infected hMDCs recapitulates known binding motifs of class I HLA alleles. Peptide sequence motif logos associated with each cluster in the optimal clustering solution determined by GibbsCluster 2.0^28,29^ (top row for each donor) and the published peptide binding sequence motif logos (obtained from the NetMHCpan 4.1 motif viewer^30^) for the corresponding HLA alleles expressed by the donor (bottom row for each donor). The optimal number of clusters was defined as the number between 1 and 6 that maximized the Kullback-Liebler distance among the sequence distributions of the clusters.^28^

**Supplementary Figure 6.**
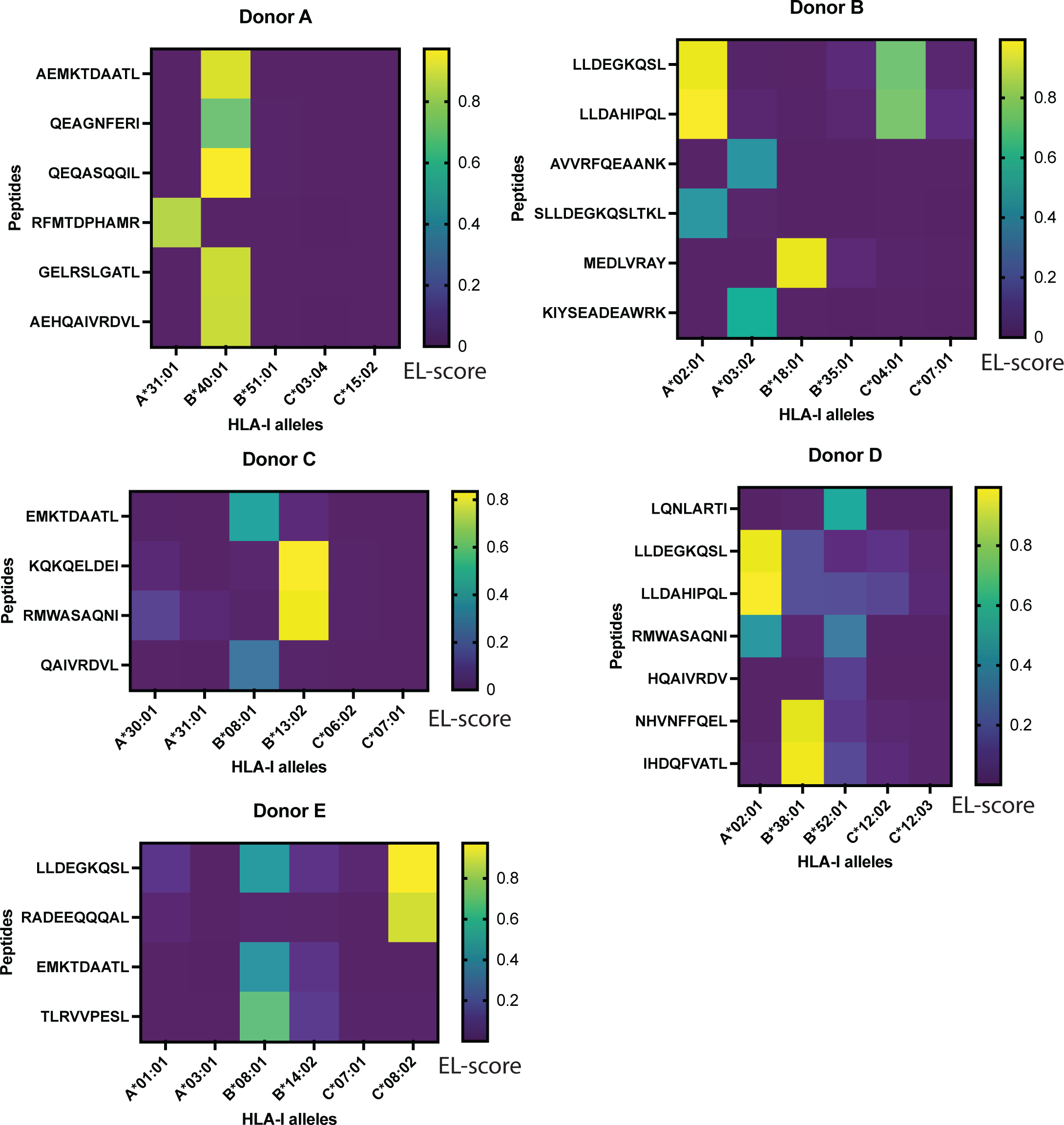
*Mtb*-derived MHC-I peptides are predicted binders of HLA alleles expressed by each donor. Heatmaps show the Eluted Ligand (EL)-score assigned to each *Mtb*-derived MHC-I peptide by NetMHCpan4.1^30^ for each class I HLA allele expressed by a given donor.

**Supplementary Figure 7.**
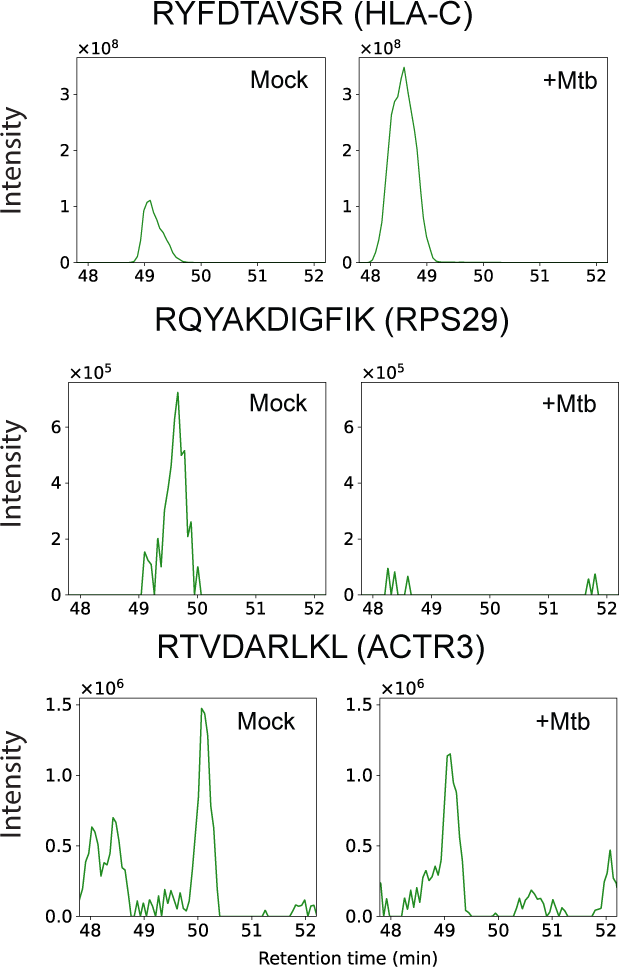
Extracted ion chromatograms of self peptides identified in DDA analyses. Representative extracted ion chromatograms of peptides identified by DDA in mock-infected hMDCs, showing MS1 signal in *Mtb-*infected and mock-infected hMDCs.

## Supplementary note 1

To maximize the ease of adoption of PathMHC, we also asked whether an analogous computational workflow implemented in a commercial proteomics analysis program could produce comparable results to our pipeline based on open-source tools. We implemented a workflow (PathMHC-PD) in Thermo Scientific’s proteomics analysis program Proteome Discoverer that can also perform alignment of precursor ion peaks across samples and select candidate peaks that are specific to the MHC repertoire of infected cells (see Methods). A workflow based on DDA alone identified 20.5 bacterial peptides on average, while PathMHC using our computational pipeline based on open-source tools identified 40 bacterial peptides on average and PathMHC-PD identified 34 bacterial peptides on average (Supplementary figure 2 a). Whereas on average 1.02% of peptides identified in DDA runs were bacterial peptides, 5.97% of peptides identified in PathMHC PRM runs using our open-source computational workflow and 8.20% of peptides identified in PathMHC-PD PRM runs were bacterial peptides (Supplementary figure 2 b). These results show that either our open-source implementation of the PathMHC computational pipeline or an analogous workflow implemented in a commercial proteomics analysis program can provide comparable performance in enriching and identifying pathogen-derived MHC peptides.

**Supplementary Table 1.**
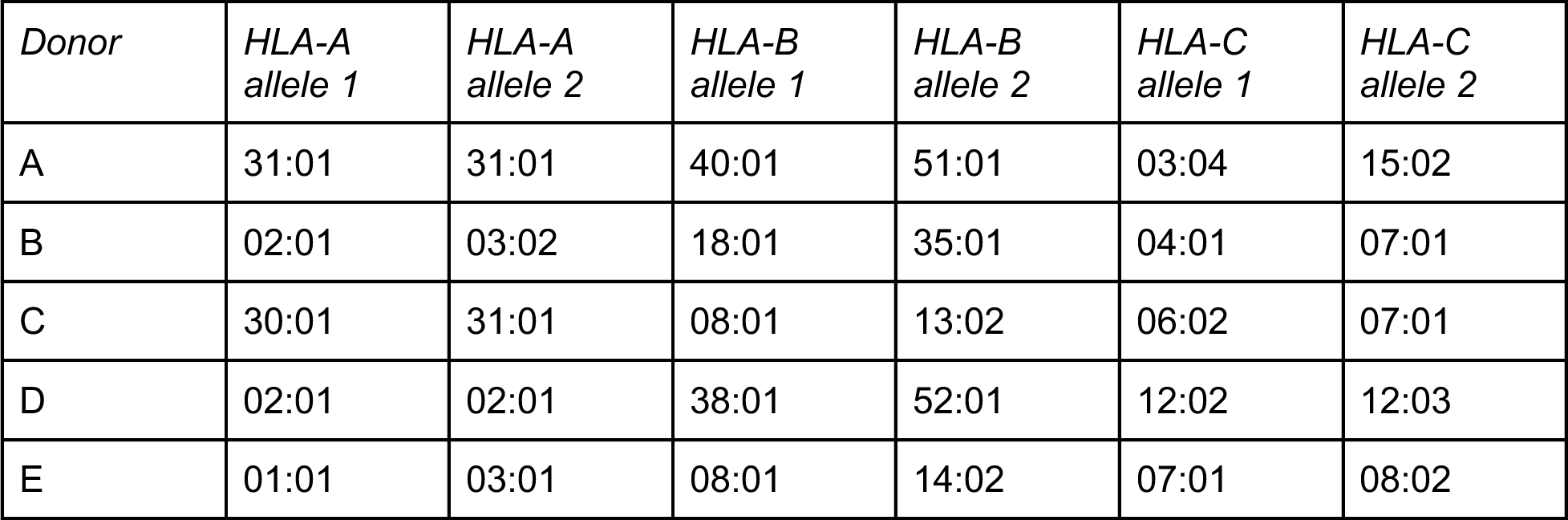
HLA types of primary cell donors.

**Supplementary Table 2.** Putative *Mtb* peptide IDs invalidated by SureQuant analyses.

| *Sequence* | *Source protein* |
| --- | --- |
| HPLLEAVVL | Rv0552 |
| EVADLPDAL | Rv2295 |
| KLADFGLVRA | PknL |
| VPVTFEGGQMPI | RpIO |

